# An octadecameric *O-*glucosyltransferase generates diversity in antibody epitopes on variant surface antigens in African trypanosomes

**DOI:** 10.64898/2026.01.27.701950

**Authors:** Qi Zhong, Joseph D Barritt, Carla Gilabert Carbajo, Lilian Magnus, Olivia Wright-Paramio, Laura Gordon, Wuxi Song, Emmanuel Nji, Anastasia Gkeka, Eleftheria Papamanoglou, Sven A. Vida, Michael A J Ferguson, Sarah L. Rouse, Erhard Hohenester, Calvin Tiengwe

## Abstract

Immune evasion in many pathogens relies on sequence variation to generate antigenic diversity. African trypanosomes use an additional strategy where *O*-glucosylation of variant surface glycoproteins (VSGs) alters antibody epitopes and influences infection outcome, but the responsible enzyme remained elusive. Here, we show that an expression-site-associated gene (ESAG3), which is co-transcribed with the active VSG, encodes an *O*-glucosyltransferase required for VSG3 *O-*glycosylation *in vivo*. ESAG3 depletion abolishes VSG3 *O-*glycosylation and recognition of live trypanosomes by an *O*-glycan-dependent monoclonal antibody, while complementation restores both. ESAG3 has strict UDP-glucose donor specificity and modifies serine/threonine residues within VSG3 peptides *in vitro*. Cryo-electron microscopy at 3.4 Ångstrom resolution reveals that ESAG3 forms a homo-octadecameric ring with C3 symmetry, an architecture not previously observed among glycosyltransferases. Structure-guided mutagenesis shows that disruption of ESAG3 inter-subunit interfaces reduces higher-order assembly, enhances substrate conversion, and alters VSG glycoform composition. Furthermore, we show that VSG3 *O*-glycan occupancy correlates with susceptibility to complement-mediated trypanolysis in an antibody-dependent manner, establishing that ESAG3-mediated *O*-glucosylation shapes both antibody specificity and the functional outcome of the host response.

## Introduction

African trypanosomes evade host immunity primarily through antigenic variation, periodically switching expression of immunodominant variant surface antigens (VSGs), with *Trypanosoma brucei* having one of the most complex antigenic diversity among pathogens^1–3^. Each trypanosome expresses a single VSG from one of 15 sub-telomeric bloodstream expression sites (BESs), selected from an archive of >2,500 genes^4–6^, enabling chronic infections^7–10^. VSG constitutes >95% of all surface protein^11^, forming a dense monolayer of 10 million molecules per cell^12^, with antigenic diversity traditionally attributed to the hypervariable surface-exposed N-terminal domain (NTD).

All VSGs are anchored to the cell membrane via glycosylphosphatidylinositol moieties, and almost all are decorated with 1-3 *N*-glycans^13,14^. The *N*-glycans support trafficking, lateral diffusion, and coat structure but have no established direct roles in immune evasion^15–18^. Broadly, VSGs are classified into Class A (dimeric, e.g., VSG2) and Class B (either monomeric or trimeric, e.g., VSG3) based on disulphide bonding patterns and structural topology^19–22^. Many class B VSGs are *O*-glycosylated within cysteine-flanked surface-exposed loops at the membrane-distal tip of the NTD. In VSG3, two neighbouring serines, S317 and S319, are *O*-glycosylated^20,23^, with S317 having a heterogeneous set of 0/1/2/3-hexoses, while S319 has a single *O-* linked glucose^20,23^. The initiating sugar at both serines is an α1-linked *O*-glucose but the identity of the additional hexoses at S317 remains unassigned^20^. Like VSG3, VSG11 and VSG615 are also *O*-glycosylated, but not the class A VSG2, suggesting *O*-glycosylation is frequent in Class B VSGs^20^ which constitute roughly half of the genomic VSG repertoire in all *T. brucei* subspecies^24^.

VSG3 *O-*glycosylation profoundly influences early antibody responses and trypanosome virulence in mouse infection models^20,23^. Trypanosomes expressing wild-type *O-*glycosylated VSG3 cause lethal infections, while parasites expressing VSG3-S317A are cleared during the first parasitaemia wave^20^, reflecting diversified antibody repertoires elicited by VSG3-WT compared to the focused, stereotyped responses elicited by VSG3-S317A^23^. Monoclonal antibodies from these infections distinguish *O-*glycosylated and non-glycosylated VSG3, demonstrating that *O-*glycosylation generates distinct epitopes within the same VSG sequence^23^. However, the enzyme(s) responsible for generating these epitopes remained unidentified.

To our knowledge, direct attachment of α-*O*-linked glucose to serine residues within a surface-exposed di-cysteine loop has been reported only for trypanosome Class B VSGs^20^. Mammalian POGLUT1 installs β-*O*-linked glucose at serine within folded epidermal growth factor repeats to assist protein folding and trafficking^25,26^, while bacterial enzymes and toxins include both β- and α-*O-*glycosyltransferases, such as the *Legionella* effector LtpM^27^, SetA and Lgt1–3^28–30^, but these glycoproteins are not exposed to the host immune system. Thus, VSG *O*-glucosylation represents a mechanistically distinct deployment of α-*O*-glucose on an extracellular eukaryotic surface antigen, and the identity of the *O*-glycosylating enzyme is of high interest.

The active *VSG* is co-transcribed with variable cohorts of expression site-associated genes (ESAGs) from a specialised RNA polymerase I polycistronic unit^4^. *ESAG6/7* encode the transferrin receptor essential for iron uptake^31,32^, and *ESAG4* encodes a flagellar-associated receptor-type adenylate cyclase^33,34^; however, most ESAGs remain functionally uncharacterised^4,35^. ESAG3 is exceptional within this family: it is among the most abundant proteins in bloodstream-form *T. brucei*^36^ and genotype-specific variations are associated with trypanosome adaptation to different mammalian sera^37^. Phylogenetic analyses show that *ESAG3* genes are restricted to kinetoplastids, with BES-associated *ESAG3* sequences distinct from core chromosome-internal or sub-telomeric genes-related to ESAG3, otherwise known as *GRESAG3* or *ESAG3*-like paralogues found only in trypanosomatids^35,38^.

In this study, we combine trypanosome genetics, glycoproteomics, *in vitro* biochemistry, and single-particle cryo-electron microscopy to show that ESAG3 is an *O*-glucosyltransferase that generates antibody epitopes on live trypanosomes. We characterise its biochemical properties, substrate specificity, and oligomeric architecture, and show how this rare *O-*linked glycosylation influences trypanosome susceptibility to complement.

## Results

### ESAG3 is a candidate glycosyltransferase

We previously identified ESAG3 in a screen for iron-regulatory factors in *T. brucei*^39^. ESAG3 is encoded within VSG expression sites, with nine intact copies distributed across the 15 BESs in the *T. brucei* 427 strain (**Table S1**)^4,35^. AlphaFold and FoldSeek structural predictions suggested that ESAG3 adopts a GT-A glycosyltransferase fold, with closest structural homology to the collagen-modifying enzyme LH3/PLOD3 (RMSD 2.40 Å over 147 Cα pairs; **Fig S1A, B**)^40,41^. Consistent with previous evidence of BES-associated *ESAG3* genes undergoing purifying selection^4^, the predicted glycosyltransferase active-site residues and predicted GT-A fold are highly conserved across all nine BES-associated ESAG3 variants (**Fig S1C-E**). Taken together, the association of ESAG3 with VSG expression sites and its predicted enzymatic function prompted us to test whether it is the elusive VSG3 *O*-glycosyltransferase.

### ESAG3 is required to generate *O-*glycan-dependent antibody epitopes on live trypanosomes

To test this hypothesis, we used the monoclonal antibody Ab250 as an *in vivo* reporter of VSG3 *O-* glycosylation^23^. Ab250 recognises an *O-*glycan-dependent epitope on surface VSG3 that strictly requires S317. Ab250 binds VSG3-WT and VSG3-S319A, but not VSG3-S317A or the S317A/S319A double mutant(VSG3-SSAA)^23^. Because individual trypanosomes display a heterogeneous population of VSG3 glycoforms at S317 (0/1/2/3 hexoses) and/or a single glucose at S319, Ab250 binding reports the net epitope occupancy of this mixed glycoform population on each cell.

We generated three independent doxycycline-inducible pan-specific ESAG3 RNAi cell lines in VSG3-expressing bloodstream-form parasites. Doxycycline induction caused robust ESAG3 depletion, confirmed by Western blotting, while total VSG3 abundance remained unchanged (**Fig. 1A**). To control for off-target effects, we generated addback cell lines expressing an RNAi-resistant 1×TY::ESAG3^RR^ that restores ESAG3 expression following RNAi induction. We observed no significant growth defect over 4 days following RNAi induction, although two RNAi lines were leaky (RNAi-2, RNAi-3), showing partial knockdown in the absence of doxycycline induction (**Fig. 1A, B**).

**Figure 1:**
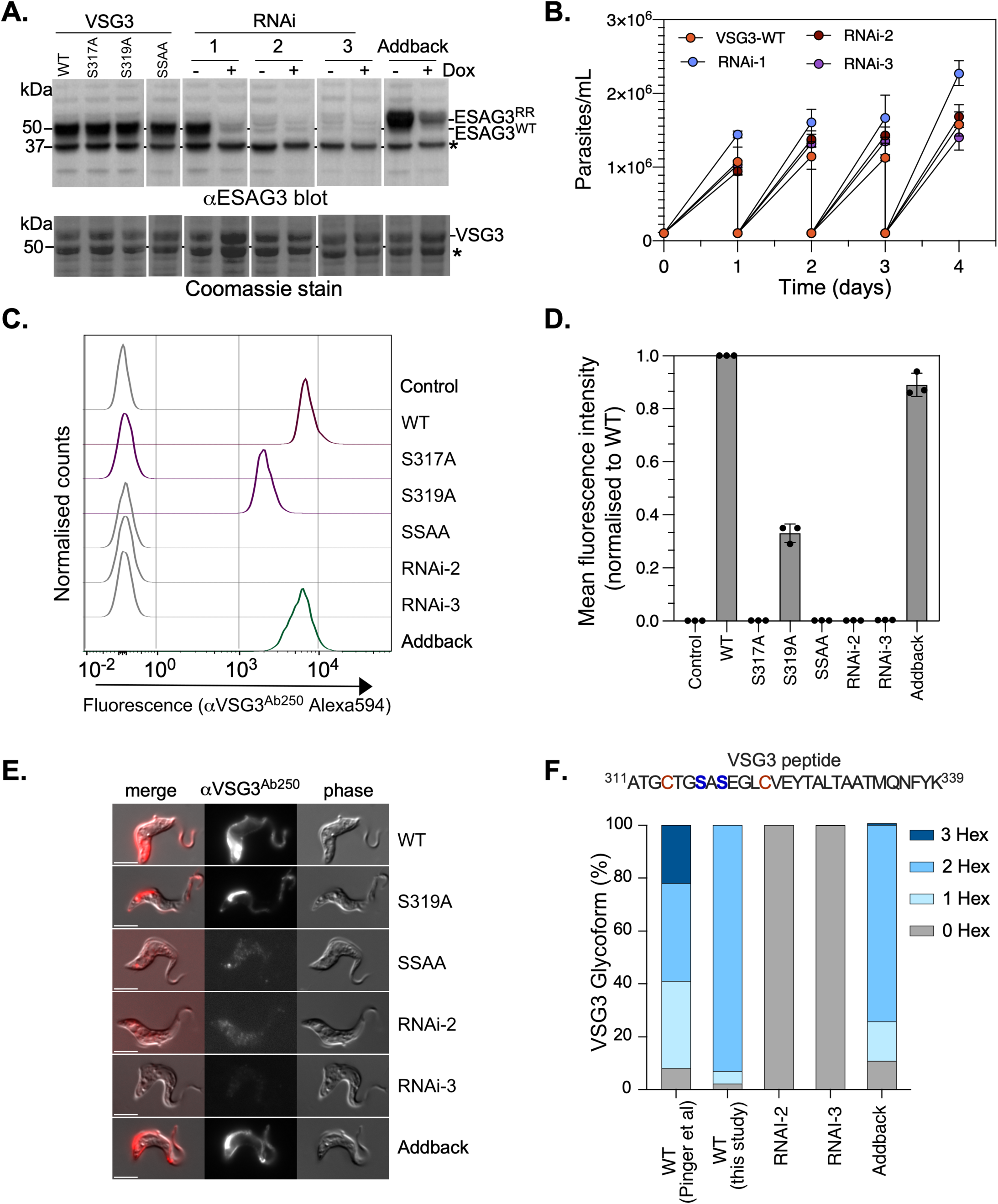
ESAG3 is required for VSG3 *O*-glucosylation *in vivo*. **A. ESAG3 depletion by RNAi and complementation**. Western blot analysis of ESAG3 expression in VSG3-expressing cells. **Top**: Anti-ESAG3 antibody detects endogenous ESAG3^WT^ and RNAi-resistant 1×TY-tagged ESAG3^RR^) in VSG3 wild-type and mutant cells (WT, S317A, S319A, SSAA), three independent ESAG3 RNAi cell lines (1, 2, 3), and addback cells, shown before (-) and after (+) 72-hour doxycycline induction. **Bottom**: Coomassie stain showing total protein with VSG3 visible at ∼50 kDa. Asterisk (*) denotes a non-specific band. Molecular weight markers (kDa) shown on left. Anti-ESAG3 was probed on two separate membranes and digitally separated for presentation. **B. Growth kinetics of ESAG3 wild type and RNAi cells.** VSG3-WT-expressing cells and three independent ESAG3 RNAi cell lines (RNAi-1, RNAi-2, RNAi-3) cultured over 4 days following doxycycline induction. Cell density was determined by haemocytometer. Data are means ± SD (n = 3 technical replicates). **C. Flow cytometry analysis of Ab250 binding.** Histogram plots showing Ab250-Alexa594 fluorescence intensity distribution of 10,000 single cells. Cells include unstained control (Control), VSG3 WT, S317A, S319A, SSAA, ESAG3 RNAi-1, RNAi-2, and addback. Cells were incubated with monoclonal antibody Ab250 conjugated to Alexa594 at 4°C for 10 minutes before analysis. RNAi-3 showed identical results to RNAi-1 and RNAi-2 and is omitted for clarity. **D. Quantification of Ab250 binding by flow cytometry.** Median fluorescence intensity (MFI) from panel C, normalised relative to VSG3 WT-expressing cells. Individual data points shown as dots for n ≥ 3 independent experiments. Data show means ± SD. RNAi quantification data (n = 9) is combination of all three cell lines. **E. ESAG3 depletion abolishes Ab250 binding to surface VSG3**. Immunofluorescence analysis of live trypanosomes incubated with Ab250 monoclonal antibody conjugated to Alexa594 (red fluorescence) at 4°C for 10 minutes. VSG3-WT, SSAA, ESAG3 RNAi-2, RNAi-3, and addback cells shown. Merge (left), αVSG3-Ab250 fluorescence (middle), and phase contrast (right) are shown. Bright posterior signal in Ab250-positive cells (WT, S319A, addback) represents VSG-antibody complexes concentrated at the flagellar pocket during forward motility. Representative images from at least three independent experiments. Scale bar = 5 μm. **F. ESAG3 depletion abolishes VSG3 *O-*glycosylation *in vivo*.** Accurate-mass extracted ion profiling of the VSG3 glycopeptide ^311^ATGCTGSASEGLCVEYTALTAATMQNFYK^339^ by LC-MS. Glycoform percentages were calculated from integrated peak areas of [M+3H]³⁺ ions corresponding to 0-, 1-, 2-, and 3-hexose species (m/z 1035.13, 1089.14, 1143.16, and 1197.18, respectively). ESAG3 RNAi lines (RNAi-2, RNAi-3) show near-complete loss of *O*-glycosylation (0Hex ≥99.96%), while addback restores the glycoform distribution toward wild-type. Wild-type (this study) values: 0Hex 2.2%, 1Hex 4.7%, 2Hex 93.1%, 3Hex 0%. Values from Pinger et al. (2018) shown for comparison. Glycosylated serine residues S317 and S319 (blue) are indicated on the peptide sequence. See **Fig. S2** for extracted ion chromatograms.

Flow cytometry analysis showed that wild-type VSG3 cells exhibited strong Ab250 binding, while S317A and SSAA mutants showed near-background signal (0.13-0.15% of wild-type) (**Fig. 1C, D**). S319A cells retained ∼32% of wild-type signal, indicating S319 contributes to recognition of the Ab250 epitope of VSG3^23^. All three ESAG3 RNAi lines reduced Ab250 binding to 0.14-0.30% of wild-type levels, phenocopying the S317A mutant. Addback cells restored Ab250 recognition to ∼87% of wild type, demonstrating that epitope loss resulted specifically from ESAG3 depletion.

Immunofluorescence microscopy confirmed the above findings (**Fig. 1E**). Wild-type VSG3 and S319A cells showed strong Ab250 surface staining, while SSAA and ESAG3 RNAi cells showed near-background signal; and addback restored staining. The intense posterior signal in Ab250-positive cells is consistent with hydrodynamic transport of VSG-antibody complexes toward the flagellar pocket^42^. Whilst it is not our aim to map the epitope, the complete loss of binding in S317A and substantial reduction in S319A establish that *O-*glycosylation at both positions are critical contributors to the Ab250-recognised epitope. The phenocopy of VSG3-S317A by ESAG3 depletion, and restoration of Ab250 binding by RNAi-resistant ESAG3, establish that ESAG3 is essential to generate the *O-*glycan-dependent antibody epitope on VSG3 *in vivo*.

### ESAG3 depletion abolishes VSG3 *O*-glycosylation *in vivo*

To directly quantify the contribution of ESAG3 to native VSG3 *O*-glycosylation, we performed accurate-mass extracted ion profiling of the VSG3 glycopeptide A311–K339 (ATGCTGSASEGLCVEYTALTAATMQNFYK) by LC-MS on purified VSG3 purified from ESAG3 RNAi, addback, and wild-type trypanosomes (**Fig. 1F; Fig. S2)**. In wild-type parasites, the 2-hexose glycoform predominated (93.1%), with minor populations of 0-hexose (2.2%) and 1-hexose (4.7%) species and no detectable 3-hexose glycoform. Both independent ESAG3 RNAi lines completely abolished VSG3 *O*-glycosylation, and expression of RNAi-resistant ESAG3 restored the glycoform distribution toward wild-type levels (0Hex 10.8%; 1Hex 14.9%; 2Hex 74.3%), confirming that glycoform loss is specifically attributable to ESAG3 depletion. Methionine-oxidised forms of the glycopeptide showed equivalent glycoform distributions, indicating that oxidation at M333 does not influence *O*-glycan occupancy. Inspection of the 1-hexose channel revealed two co-eluting species in both wild-type (3.5% and 1.2%) and addback (11.1% and 3.8%) parasites, consistent with ESAG3 adding glucose to both S317 and S319 as distinct mono-glucosylated isoforms, with preference for S317 as the primary modification site^20,23^. A single 2-hexose species was detected for all glycosylated samples. Together, these data establish that ESAG3 is essential for generating heterogeneous VSG3 *O*-glycoforms *in vivo*.

### ESAG3 is a Mn²⁺-dependent *O*-glucosyltransferase

Having established that ESAG3 is essential for generating *O-*glycan-dependent antibody epitopes (**Fig. 1**), we next sought to define the glycosyltransferase activity and nucleotide-sugar donor specificity. Prior studies identified the initiating sugar at both S317 and S319 as α1-linked *O*-glucose^20,23^, leading us to test whether ESAG3 functions as an *O*-glucosyltransferase *in vitro*. We expressed and affinity-purified recombinant ESAG3 from culture supernatants of Expi293F cells (**Fig. 2A**).

**Figure 2.**
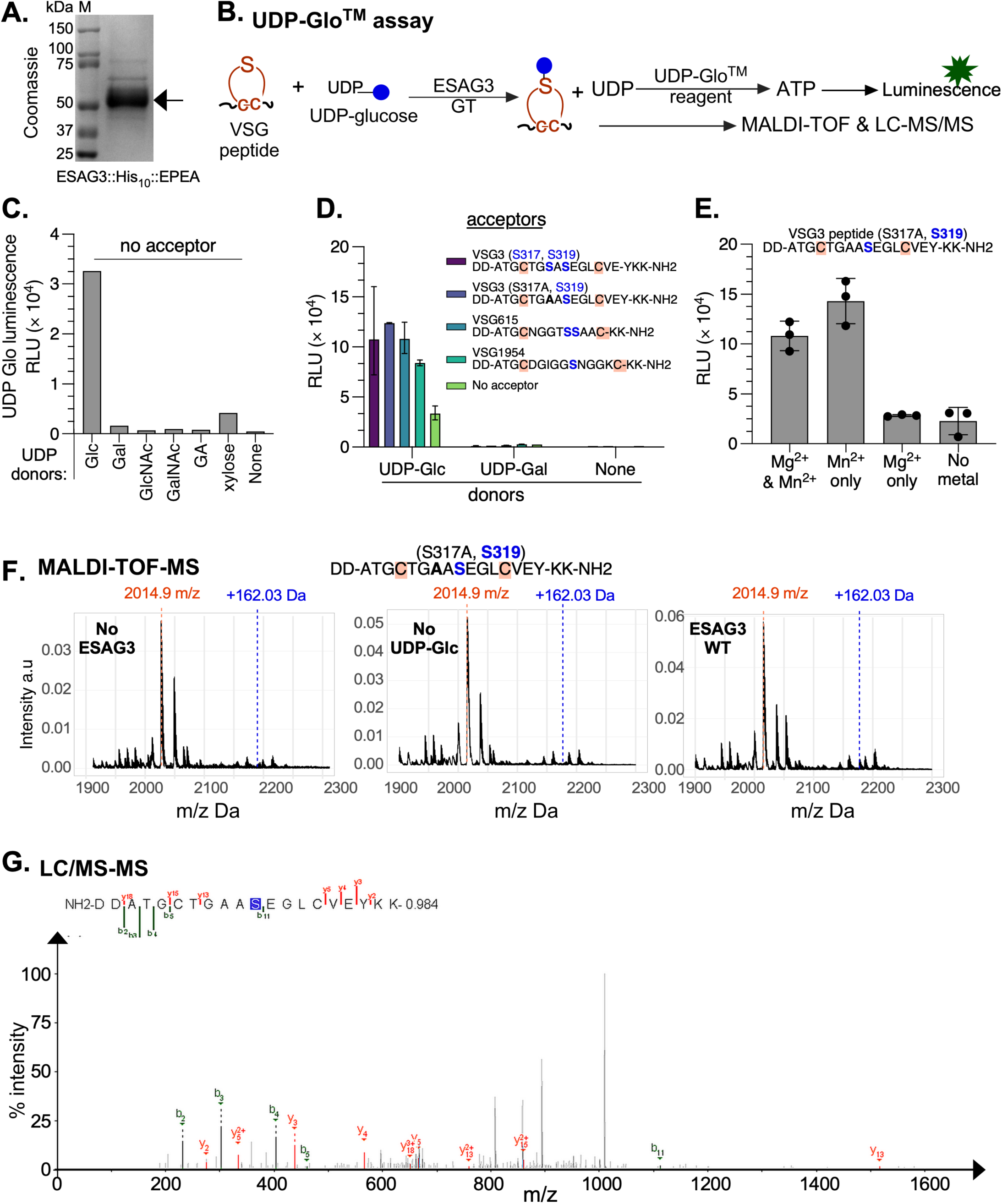
ESAG3 is a Mn²⁺-dependent O-glucosyltransferase with strict UDP-glucose donor specificity. **A. Purification**. Coomassie-stained SDS-PAGE of affinity-purified recombinant ESAG3 with C-terminal His₁₀-EPEA tag) expressed and purified from Expi293F cells. Arrow indicates ESAG3 monomer (∼50 kDa). M, molecular weight markers. **B. UDP-Glo™ assay schematic.** ESAG3-catalyzed glucosylation releases UDP from UDP-glucose, which is converted to ATP by UDP-Glo™ reagent, generating luminescence proportional to glycosyltransferase activity. Parallel mass spectrometry validates glucose addition. **C. UDP-sugar donor specificity screen without acceptor.** Purified ESAG3 was incubated with indicated UDP-sugar donors and UDP release quantified by luminescence. ESAG3 hydrolyses UDP-glucose specifically (68-fold above background), while UDP-galactose (Gal), UDP-GlcNAc, UDP-GalNAc, and UDP-glucuronic acid (GA) shows negligible activity. Single experiment; RLU, relative light units. **D. VSG peptide acceptor specificity.** ESAG3 was incubated with indicated UDP-sugar donors and synthetic VSG peptides (sequences shown; putative *O*-glucosylation sites Ser in blue). With UDP-glucose, all VSG peptides show 2.7–4-fold enhanced activity over no-acceptor baseline. UDP-galactose showed negligible activity. Y-axis: UDP-Glo luminescence (RLU × 10⁴). Bars show mean ± range (n=2 technical replicates). **E. Metal cofactor requirement.** ESAG3 activity with VSG3 (S317A,S319) peptide and UDP-glucose requires Mn²⁺. Mn²⁺ alone or combined with Mg²⁺ shows optimal activity; Mg²⁺ alone or omission of divalent cations shows only baseline activity. Bars show mean with individual replicates overlaid (n=3). **F. MALDI-TOF MS confirms glucosylation.** Analysis of VSG3 (S317A,S319) peptide DD-ATGCTGAASEGLCVEY-KK-NH₂ (expected mass 2014.9 m/z with disulfide bond). **Left:** Peptide-only control, no ESAG3. **Middle:** Reaction lacking UDP-glucose. **Right:** Complete reaction generates glucosylated product with +162.03 Da mass increase (blue annotation), consistent with hexose addition. Unmodified peptide indicated by red dotted line. Site-specific modification confirmed by LC-MS/MS (**Fig. 2G and S3)**. **G. LC-MS/MS confirms site-specific glucosylation at S319.** Representative LC-MS/MS spectrum showing +162.05 Da modification on VSG3 (S317A,S319) peptide. Annotated fragment ions (b-ions in green, y-ions in orange) localise hexose to serine 11 of the peptide sequence (S11, corresponding to S319 in full-length VSG3). Site localisation is supported by the y₁₃ ion series (singly and doubly charged) that retains the +162 Da mass shift. This modification was detected only in samples containing ESAG3 and UDP-glucose. Additional modification at threonine 7 (T7, corresponding to T315 in full-length VSG3) was also detected across replicates, with alternative site assignments detailed in **Fig. S3**. Expected mass accounts for C-terminal amidation (–0.984 Da) from peptide synthesis.

To determine nucleotide-sugar donor specificity, we used the UDP-Glo™ bioluminescence assay, which converts released UDP into luminescent signal proportional to activity^43,44^ (**Fig. 2B**). Following established methodology for UDP-sugar donor screening without acceptor substrates^44^, ESAG3 showed robust UDP release exclusively with UDP-glucose, whilst with UDP-galactose, UDP-*N*-acetylglucosamine, UDP-*N*-acetylgalactosamine, UDP-glucuronic acid, and UDP-xylose, it showed negligible activity (**Fig. 2C**). The absence of detectable hydrolysis with UDP-galactose, the C4-epimer of UDP-glucose, indicates strict donor selectivity for UDP-glucose.

To test whether VSG peptides function as sugar acceptor substrates, we used peptides containing the native di-cysteine loops surrounding known *O*-glycosylation sites. For VSG3, we used residues 311-326: DD-_311_ATGCTG**S**A**S**EGLCVEY_326_-KK) flanked by charge-complementary DD/KK sequences to enhance solubility. To simplify site-specific analysis, we used a VSG3 S317A mutant peptide retaining only S319 (ATGCTG**A**A**S**EGLCVEY). Using UDP-glucose as donor, ESAG3 showed enhanced activity, above no-acceptor baseline, for all VSG peptides (VSG3 (S317, S319), VSG3 (S317A, S319), VSG615 and VSG1954) tested (**Fig. 2D**), suggesting multiple Class B VSG sequences stimulate UDP release. No comparable activity was detected with UDP-galactose as donor, consistent with the UDP-glucose specificity established above.

ESAG3 activity required Mn²⁺, with Mg²⁺ alone showing significantly reduced activity (**Fig. 2E**), consistent with Mn²⁺-dependence characteristic of GT-A fold glycosyltransferases and distinguishing ESAG3 from related Mg²⁺-dependent enzymes^45^.

To confirm site-specific glucosylation, we analysed ESAG3-treated VSG3 (S317A, S319) peptides by mass spectrometry. MALDI-TOF MS detected a +162.03 Da mass product only in complete reactions containing both ESAG3 and UDP-glucose (**Fig. 2F**), while LC-MS/MS confirmed glucosylated peptide under these conditions (**Fig. 2G, Fig. S3)**. LC/MS-MS localised glucosylation to T315 and S319 residues within the conserved cysteine-flanked loop (**Fig. 2G, Fig. S3)**. Collectively, these data establish ESAG3 as a Mn²⁺-dependent *O-*glucosyltransferase with strict UDP-glucose specificity and demonstrate direct glucosylation of the known VSG3 peptide substrate *in vitro*.

### ESAG3 assembles as an octadecameric ring with hierarchical organisation

Glycosyltransferases can function as monomers or oligomeric assemblies^45^. To define the oligomeric state of ESAG3, we analysed purified recombinant protein by size-exclusion chromatography. ESAG3 eluted predominantly as a large complex (>669 kDa), with a minor peak at ∼50 kDa (**Fig. 3A, B**). To confirm this oligomeric state represents the native form, we compared recombinant ESAG3 with endogenous ESAG3 in whole-cell lysates from wild-type trypanosomes by blue native PAGE and anti-ESAG3 Western blotting. Both recombinant and native ESAG3 comigrated as an ∼720-1,024 kDa complex, with minor lower molecular weight oligomeric species visible in the 66-242 kDa range (**Fig. 3C**). These data indicate that recombinant ESAG3 recapitulates the native oligomeric assembly, supporting the physiological relevance of the recombinant complex.

**Figure 3:**
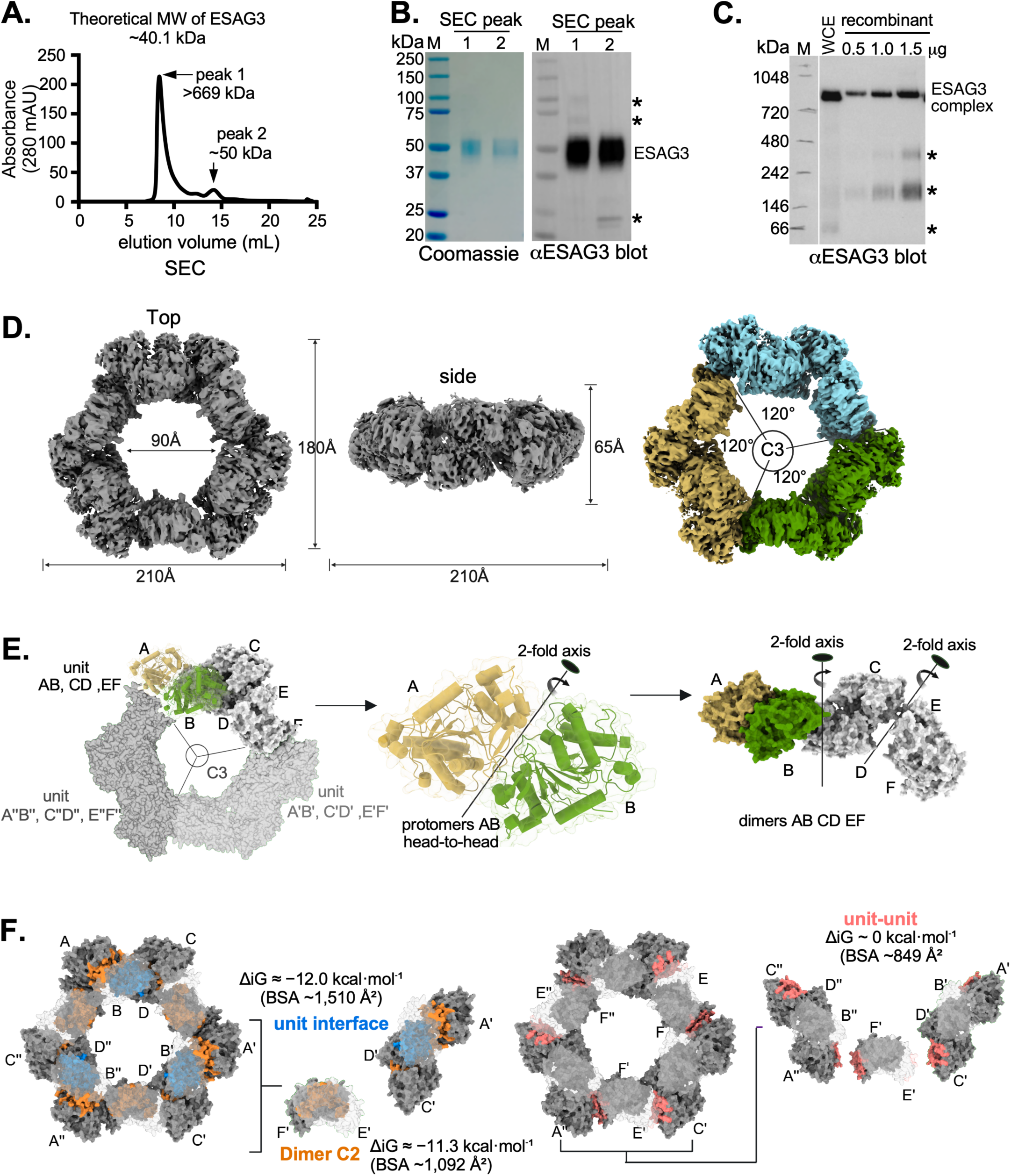
ESAG3 forms an unprecedented 18-protomer ring with hierarchical assembly. **A. ESAG3 oligomeric state by size-exclusion chromatography (SEC).** Recombinant ESAG3 (residues 23-368, C-terminal His₁₀-EPEA tag) expressed in Expi293F cells and purified from culture supernatants by immobilized metal affinity chromatography. SEC shows a major peak at >669 kDa and minor peak at ∼50 kDa (theoretical monomer 40.1 kDa). **B. SEC fraction analysis.** Coomassie staining and anti-ESAG3 Western blot of peak fractions from panel A. Arrow indicates ESAG3; asterisks mark bands recognized by anti-ESAG3 antibody. M, molecular weight markers **C. Recombinant ESAG3 comigrates with endogenous ESAG3 from whole cell trypanosome extracts.** Blue native PAGE and anti-ESAG3 Western blot of purified recombinant ESAG3 from peak 1 in (**A**) loaded at 0.5, 1.0, and 1.5 µg alongside whole cell extracts (WCE) from wild-type trypanosomes. Both WCE and recombinant ESAG3 migrate at ∼720–1,048 kDa, with minor lower MW species indicated by asterisks. M, molecular weight markers **D. Cryo-EM structure of the ESAG3 octadecamer.** Top and side views (**left, middle**) show closed toroidal architecture with dimensions indicated. **Right:** C3-symmetric organisation showing three repeating six-protomer units (green/gold, dark blue/light blue, and dark grey/light grey) forming the 18-protomer ring. The overall structure was refined to 3.4 Å resolution; and the six-protomer unit refined to 3.0 Å. See **Fig. S4** for reconstruction workflow. **E. Schematic representation of ESAG3 assembly.** Within each six-protomer unit, protomers form antiparallel head-to-head dimers via 2-fold rotational symmetry: dimer AB (gold/green), dimer CD, and dimer EF. These three dimers assemble to form each six-protomer unit. Three such units related by C3 symmetry (120° rotations) complete the 18-protomer ring. Symmetry axes indicated. **F. Interface energetics calculated by PDBePISA. Left:** Each dimer interfaces (orange, nine total: AB, CD, EF in each unit) buries ∼1,092 Å² per interface with ΔiG = −11.3 kcal·mol⁻¹. **Middle:** Each six-protomer unit stabilisation interface (blue) buries ∼1,510 Å² per interface with ΔiG = −12.0 kcal·mol⁻¹. **Right:** Each ring-closure interface (pink, six contacts between units; examples shown: Eʹ–Cʹ, Fʹ–Aʹʹ) bury ∼849 Å² per interface with ΔiG ≈ 0 kcal·mol⁻¹. BSA, buried surface area; ΔiG, interface free energy.

Given the higher-order oligomeric state of native ESAG3 and the absence of an experimental structure for a kinetoplastid UDP-dependent glycosyltransferase, we determined the structure of purified ESAG3 by single-particle cryo-electron microscopy. 2D classification revealed ring-shaped particles, and 3D reconstruction yielded a 3.4 Å map of a homo-octadecameric complex which adopted a closed toroidal architecture of ∼210 Å outer diameter with ∼90 Å central cavity, 65 Å height, and comprised of three repeating six-protomer units related by C3 symmetry (**Fig. 3D; Fig. S4; Table S2**). Symmetry expansion followed by focused refinement of one six-protomer unit further improved the local resolution to 3.0 Å. We then fitted an AlphaFold3-predicted monomeric model into the cryo-EM density as an initial template, then iteratively rebuilt and refined it against the experimental map (**Fig. S4**). Because AlphaFold3 could not predict the higher order hexameric or octadecameric assembly, we generated the full oligomeric assembly by applying symmetry operations derived from the reconstruction.

The 18 protomers organise hierarchically through nested symmetry elements (**Fig. 3E**). Within each six-protomer unit, protomers associate as antiparallel head-to-head dimers through 2-fold rotational axes such that protomers A and B form dimer AB, C and D form dimer CD, and E and F form dimer EF. These three dimers then assemble to complete each six-protomer unit. Three such units, related by 120° rotations about the C3 axis, form the octadecameric ring.

Interface analysis by PDBePISA estimated distinct interface properties at each organisational level (**Fig. 3F**). Each of the nine head-to-head dimer interfaces (AB, CD, EF in each of three units) buries ∼1,092 Å² per interface with favourable interaction energy (ΔiG = −11.3 kcal·mol⁻¹), consistent with dimers representing a major stabilising interaction within the assembly. Interfaces stabilising each six-protomer unit bury ∼1,510 Å² per interface with ΔiG = −12.0 kcal·mol⁻¹. In contrast, contacts between six-protomer units that close the ring (six interfaces total; e.g., Eʹ–Cʹ and Fʹ–Aʹʹ) bury only ∼849 Å² per interface with near-zero interaction energy (ΔiG ≈ 0 kcal·mol⁻¹). This interface hierarchy indicates that dimers and six-protomer units are consistent with relatively stable assembly intermediates, while ring closure is mediated by comparatively weak inter-unit contacts.

Structural similarity searches failed to identify a comparable oligomeric assembly among known glycosyltransferases or other PDB entries, establishing the octadecameric architecture with C3 symmetry as unprecedented. This atypical oligomeric glycosyltransferase architecture provided a structural framework for probing ESAG3 catalytic mechanism and interface properties by targeted mutagenesis.

### ESAG3 adopts a canonical GT-A fold with a conserved active site

Glycosyltransferases of the GT-A family are characterised by a Rossmann-like fold containing a metal–ion-coordinating canonical DXD motif essential for catalysis^45^. To characterise the active site, we aligned ESAG3 with the LH3/PLOD3 (LH3) experimental structure. As expected, ESAG3 superposition with LH3 GT domain revealed close structural similarity (RMSD 2.40 Å over 147 Cα pairs), consistent with ESAG3 adopting the canonical GT-A fold (**Fig. 4A, S5**). All hallmark GT-A catalytic features aligned precisely: the metal-binding motif (D110–D113) and C-terminal His (H297) superimposed with LH3 D112–D115 and H253, respectively; the glycine-rich loop (G215–G216) overlaid with LH3 G167–G168; and a conserved acidic-polar pair (E244/Q245) aligned with LH3 D191/Q192. The spatial positioning of UDP and Mn²⁺ from the LH3 structure mapped directly onto the equivalent binding pocket in ESAG3, suggesting conservation of the nucleotide-sugar binding site. All catalytic residues are contained within individual protomers, indicating the GT-A catalytic machinery is complete in monomeric units (**Fig. S6A**).

**Figure 4.**
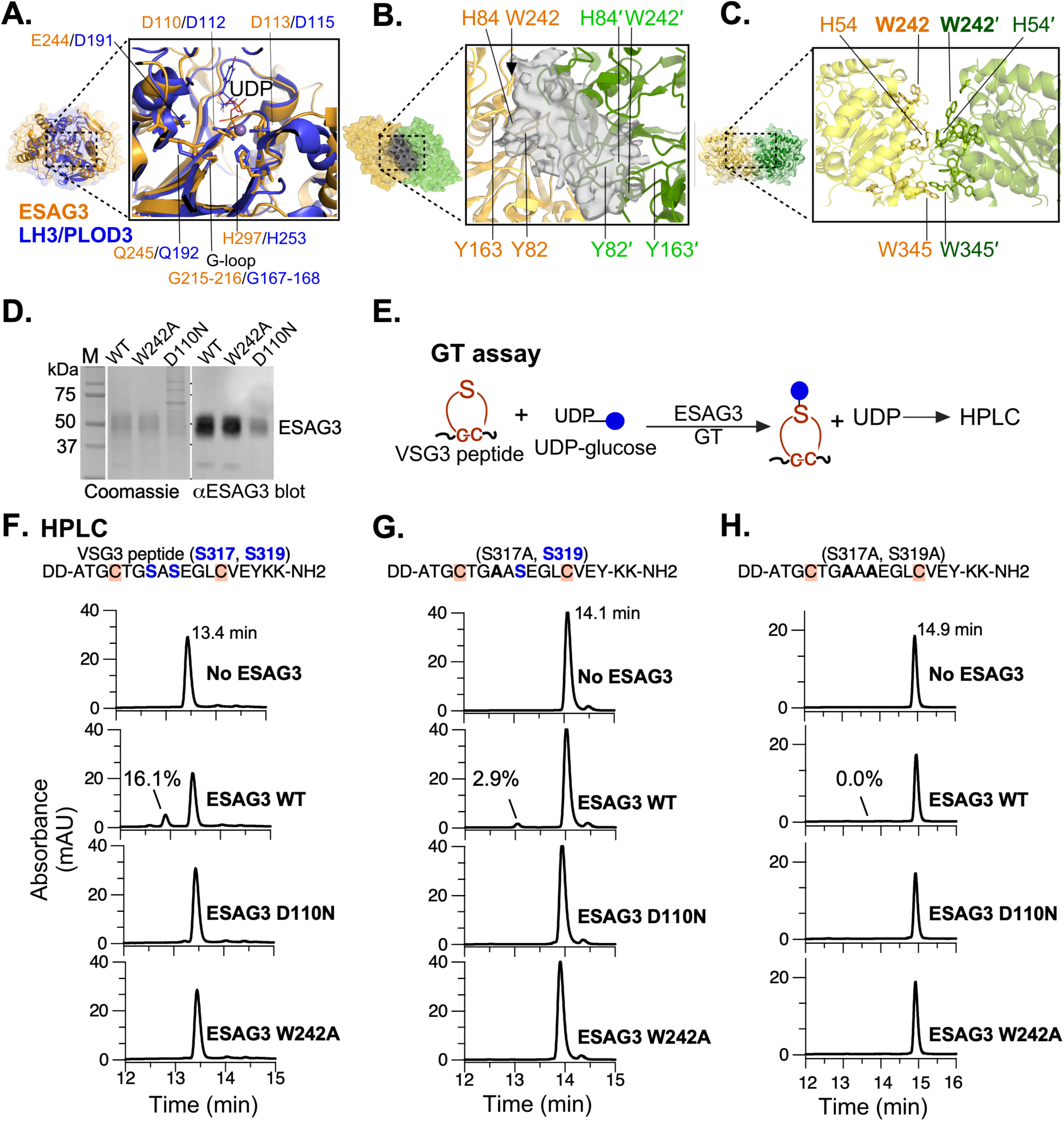
ESAG3 adopts a GT-A fold with conserved active site essential for catalysis. **A. Structural alignment with human LH3/PLOD3. Left:** Superposition of ESAG3 (sand) with LH3 GT domain (blue, PDB: 6FXR) showing conserved GT-A architecture. **Right**: Active site close-up showing conservation of catalytic residues including metal-coordinating DXXD motif (D110/D113, LH3 D112/D115) and C-terminal His (H297, LH3 H253), glycine-rich loop (G215-216, LH3 G167-168), and an acidic-polar pair (E244/Q245, LH3 D191/Q192). Mn²⁺ (purple sphere) and UDP (sticks) from LH3 crystal structure (PDB: 6FXR) indicate conserved nucleotide-sugar binding pocket position. **B. Dimer interface organisation.** Two protomers (yellow, green) assembled head-to-head showing active-site clefts facing the interface. Unassigned density (grey mesh) observed in outward-facing dimers. Aromatic residues from both protomers (H84/H84ʹ, W242/W242ʹ, Y82/Y82ʹ, Y163/Y163ʹ) line the interface cavity. Dashed boxes indicate regions shown in panel C. **C. Close-up of aromatic-rich interface.** H54/H54ʹ and W345/W345ʹ positioned at contact distance across the dimer interface. Black arrows indicate residues selected for mutagenesis to test interface contribution to assembly (see **Fig. S6** for enlarged view). **D.** Recombinant ESAG3 wild-type and mutants (W242A, D110N) were expressed and purified, confirmed by Coomassie-stained SDS-PAGE (left) and anti-ESAG3 western blot (right). **E. Glucosyltransferase assay schematic.** VSG3 peptides containing cysteine-flanked loop with disulfide bridge were incubated with UDP-glucose, Mn²⁺, and purified ESAG3. Products analysed by HPLC (280 nm detection). **F-H. HPLC analysis of VSG3 peptide variants** incubated with no enzyme (control), wild-type ESAG3 or the catalytic mutants (D110N and W242A). **F.** Wild type ESAG3 converted the VSG3 (S317, S319) peptide to product (16.1% conversion); D110N and W242A show no detectable product (0%). **G.** The VSG3 (S317A, S319) single mutant peptide retains activity with wild-type ESAG3 (2.9% substrate conversion), confirming S319 can serve as acceptor; D110N and W242A shows no product. **H.** The VSG3 (S317A, S319A) double-mutant peptide shows no product formation with wild-type ESAG3 or either catalytic mutant.

Examination of the dimer interface revealed that when two protomers assemble head-to-head (e.g., protomers A and B forming dimer AB), their active-site clefts face each other across the interface (**Fig. 4B, Fig. S6**). While each protomer contains a complete GT-A catalytic core, this arrangement creates an enlarged inter-protomer cavity. In the octadecameric ring, three of the nine dimers present outward-facing, solvent-accessible pockets, while the remaining six are oriented above or below the plane of the ring and are partially occluded (**Fig S7**). Within the three outward-facing dimers, we observed additional density in the inter-protomer cavity (**Fig. 4B**). The identity of this density remains undefined and could not be reliably modelled. The cavity is lined by aromatic residues from both protomers (H84/H84ʹ, Y82/Y82ʹ, W242/W242ʹ, Y163/Y163ʹ), with H54/H54ʹ and W345/W345ʹ positioned directly opposite one another at the interface and predicted to form π–π stacking interactions (**Fig. 4B, C, S6B**).

To test the functional importance of conserved active-site residues, we generated D110N and W242A mutants. D110 is the first aspartate of the metal-coordinating DXXD motif essential for GT-A catalysis, while W242 lines the UDP-binding pocket and likely acts as a gating residues controlling substrate access (**Fig. 4B, C**). First, we expressed recombinant wild-type and ESAG3 mutants and confirmed their purity by SDS-PAGE and anti-ESAG3 immunoblotting (**Fig. 4D**). Next, we used HPLC analysis of VSG3 peptides to determine substrate glucosylation (**Fig. 4E-H**). Wild-type ESAG3 converted VSG3 (S317, S319) peptide to an earlier-eluting product (16.1% conversion) that was assigned as the mono-glucosylated form (confirmed below for a hyperactive ESAG3 mutant, see Fig. 5). Both D110N and W242A completely abolished detectable activity, showing only the unmodified substrate peak with no detectable product (0% conversion; **Fig. 4F**). VSG3 (S317A, S319) peptide retained activity with wild-type ESAG3 but with substantially reduced conversion (2.9% versus 16.1% for the parental peptide, ∼18% of wild type), demonstrating that S319 alone can serve as an acceptor but conversion is strongly reduced in the absence of S317 (**Fig. 4G**). D110N and W242A again showed no detectable product formation (**Fig. 4G**). The VSG3 (S317A, S319A) double mutant peptide showed no product formation with wild type ESAG3 or either catalytic mutant (**Fig. 4H**), demonstrating that removal of both serines abolishes detectable activity in this assay. Collectively, these results identify D110 and W242 as specifically required for ESAG3 catalytic activity.

**Figure 5.**
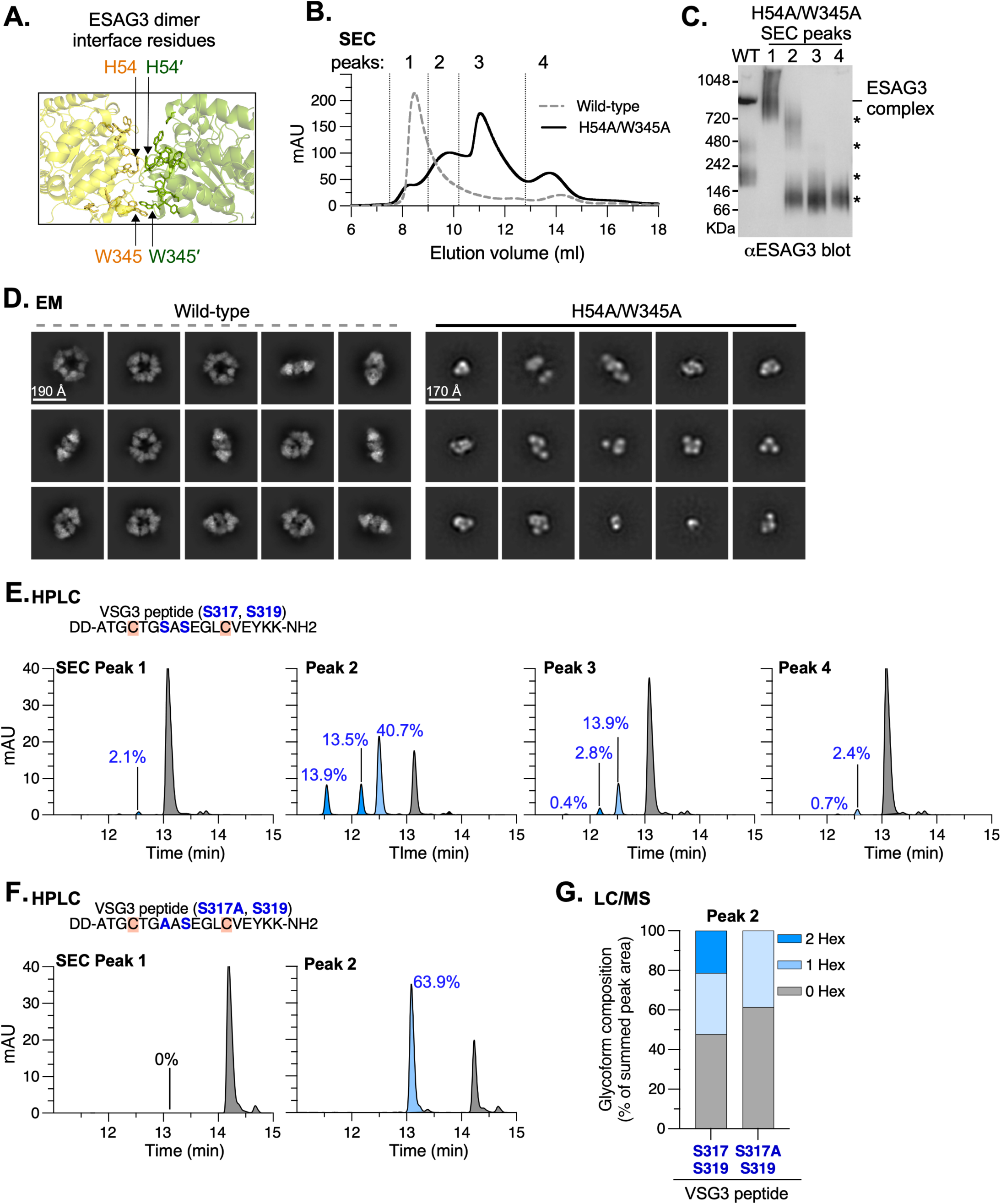
An ESAG3 inter-subunit interface mutant disrupts octadecamer assembly and alters VSG3 peptide glucosylation *in vitro*. **A.** Close-up of the ESAG3 dimer interface showing H54/H54ʹ and W345/W345ʹ (yellow and green) from adjacent ESAG3 protomers. H54 and W345 were targeted for alanine substitution. **B. Size-exclusion chromatography (SEC) of wild-type (grey dashed) and H54A/W345A (black) ESAG3**. Wild-type elutes predominantly as a high-molecular-weight complex; H54A/W345A resolves into four peak fractions (1–4). **C. Blue native PAGE and anti-ESAG3 Western blot of wild-type ESAG3 and the four H54A/W345A SEC fractions from (B).** Wild-type ESAG3 (WT) migrates predominantly as a single high-molecular-weight complex; the H54A/W345A fractions are enriched as lower-molecular weights species (asterisks). **D.** Representative 2D class averages of wild-type (from cryo-EM) and H54A/W345A ESAG3 from negative-stain EM. Wild-type particles are predominantly ring-shaped; H54A/W345A forms smaller, disrupted particles. **E-F.** Reverse-phase HPLC analysis of reactions containing VSG3 (S317, S319) peptide with H54A/W345A SEC peak fraction 1 or peak fraction 2. Apparent conversion is indicated for each resolved product peak. **F.** As in (E), using VSG3 (S317A, S319) peptide. **G.** LC-MS quantification of reaction products generated by H54A/W345A peak fraction 2 from VSG3 (S317, S319) or VSG3 (S317A, S319) peptides. Relative glycoform composition (% of summed peak area) is shown for unmodified (0 Hex), monohexosylated (1 Hex) and dihexosylated (2 Hex) species. No dihexosylated product was detected with the S317A peptide.

### An ESAG3 inter-subunit interface mutant disrupts octadecamer assembly and alters VSG3 peptide glucosylation

To test whether inter-subunit contacts influence ESAG3 assembly and activity, we generated an H54A/W345A double mutant targeting residues at the dimer interface, approximately 13–23 Å from the catalytic DXXD motif (**Fig. 5A, S6**). While wild-type ESAG3 eluted predominantly as a high-molecular-weight complex, H54A/W345A resolved into four peak fractions by size-exclusion chromatography (**Fig. 5B**). To further confirm disassembly of the octadecameric ring architecture, we performed blue native PAGE and negative-stain EM, which showed progressively smaller species relative to the wild-type octadecamer (**Fig. 5C, D**).

Next, we assessed whether these lower molecular species were catalytically active. HPLC analysis of VSG3 (S317, S319) peptide of the four SEC fractions revealed substantial differences in activity: peaks 1, 3 and 4 converted 2.1%, 17.1% and 3.1% of substrate, respectively, while peak 2 was markedly more active, converting 68.1% of substrate to at least three distinguishable glucosylated products (13.9%, 13.5% and 40.7% of total peak area; **Fig. 5E,F**). Given its substantially higher conversion, we selected peak 2 for LC-MS-based extracted-ion chromatogram profiling of the reaction products (**Fig. 5G, S8**). With the VSG3 (S317, S319) peptide, unmodified, monoglucosylated and diglucosylated species represented 47.7%, 30.9% and 21.5% of the summed extracted-ion signal, respectively. By contrast, the VSG3 (S317A, S319) peptide generated only unmodified and monoglucosylated species (61.3% and 38.7%), with no detectable diglucosylated product. These data are consistent with S317 and S319 providing two acceptor sites for ESAG3-mediated glucosylation, such that mutation of S317 prevents formation of the doubly glucosylated product.

### ESAG3 is an *N-*glycosylated ER-resident protein

For ESAG3 to modify VSG3, the two proteins must encounter one another during VSG transport through the secretory pathway, but ESAG3 lacks sequence features that predict its subcellular localisation. The cryo-EM reconstruction revealed additional densities consistent with *N-*glycosylation, including well-resolved densities extending from N99 on symmetry-related protomers into the central cavity of the octadecameric ring (**Fig. 6A**). ESAG3 contains six predicted *N*-glycosylation sites based on trypanosome oligosaccharyltransferase substrate preferences, with N31, N32, N35, and N99 predicted as TbSTT3A and N39 and N315 predicted TbSTT3B substrates^14,46^.

**Figure 6.**
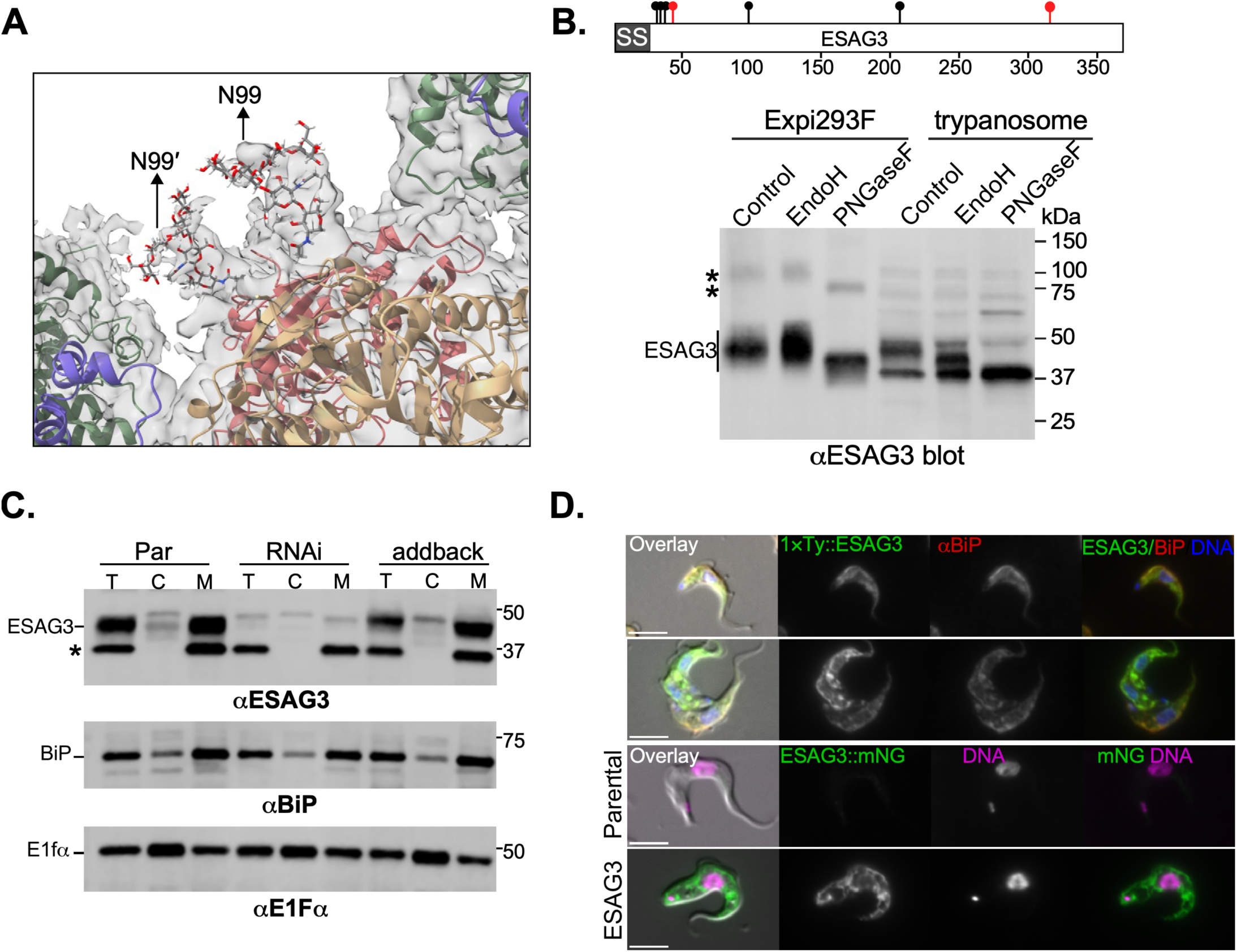
ESAG3 is an *N*-glycosylated ER-resident glycosyltransferase. **A. *N*-glycan densities in cryo-EM reconstruction.** Focused view of the ESAG3 octadecamer showing well-resolved *N*-glycan densities (red, stick representation) extending from N99 on two symmetry-related protomers (coloured surfaces), projecting into the central cavity. This represents a subset of the 18 protomers; additional *N*-glycans at N99 and other sites (N31, N32, N35, N39, N315) may be present but unresolved due to conformational flexibility. Full octadecamer structure shown in **Fig. 3**. **B. ESAG3 *N-*glycosylation analysis. Top:** Schematic of native trypanosome ESAG3 showing signal sequence (SS, grey), predicted *N*-glycosylation sites (lollipops), and molecular weight (40 kDa mature protein). Black lollipops indicate predicted complex/paucimannose-type sites (N31, N32, N35, N99; TbSTT3A substrates); red lollipops indicate predicted oligomannose-type sites (N39, N315; TbSTT3B substrates). **Bottom:** Western blot (anti-ESAG3) comparing Expi293F and trypanosome ESAG3. Samples: untreated (Control), Endoglycosidase H (EndoH), or PNGase F (PNGaseF). Expi293F ESAG3 is EndoH-resistant (complex glycans only), PNGase F-sensitive (collapses to ∼43 kDa). Trypanosome ESAG3 is both EndoH- and PNGase F-sensitive (mixed oligomannose and complex/paucimannose glycans). Asterisks indicate higher molecular weight glycosylated forms. **C. ESAG3 co-fractionates with ER marker BiP.** Bloodstream-form trypanosomes hypotonically lysed and fractionated into total (T), cytosolic (C), and membrane (M) fractions. Samples: parental cells (Par), ESAG3 RNAi (RNAi), and RNAi cells with N-terminally TY-tagged, RNAi-resistant ESAG3 addback (addback; 1×TY::ESAG3^RR^ retains native C-terminal sequence). Western blots probed with anti-ESAG3, anti-BiP (ER marker), and anti-EF1α (cytosolic marker). Both wild-type and tagged ESAG3 fractionate exclusively with BiP in membranes. Asterisk indicates non-specific band. Representative of three experiments. **D. ESAG3 localises to the endoplasmic reticulum. Top two rows:** Immunofluorescence of N-terminally tagged 1×TY::ESAG3^RR^ (anti-TY antibody, green) colocalises with BiP (anti-BiP antibody, red). DNA stained with DAPI (blue). **Bottom two rows:** Live-cell epifluorescence microscopy of parental cells (wild-type control, no fluorescence) and C-terminally tagged ESAG3::mNeonGreen (ESAG3::mNG fusion retains native signal sequence and ER retention signal) showing characteristic ER reticular pattern (green). DNA stained with Hoechst 33342 (magenta). Scale bars = 5 μm. The bottom panels show live-cell imaging of ESAG3::mNeonGreen without BiP co-staining.

To validate *N*-glycosylation, we compared recombinant ESAG3 expressed in Expi293F cells with endogenous trypanosome ESAG3 using glycosidase sensitivity assays (**Fig. 6B**). Both proteins migrated as diffuse bands consistent with heterogeneous *N-*glycan occupancy and/or processing. Endoglycosidase H treatment, which cleaves high-mannose and hybrid but not complex *N-*glycans, left Expi293F ESAG3 largely resistant, while trypanosome ESAG3 shifted to faster-migrating species, confirming Endo-H sensitive oligomannose *N-*glycan content. PNGase F treatment, which removes all *N-*glycan types, collapsed both proteins to ∼40 kDa, matching the predicted molecular weight of mature ESAG3. These results establish that ESAG3 is *N-*glycosylated and therefore enters the secretory pathway.

We next determined ESAG3 subcellular localisation by biochemical fractionation and fluorescence microscopy. Wild-type ESAG3 and N-terminally tagged 1×TY::ESAG3^RR^ co-fractionated exclusively with the ER chaperone BiP in membrane fractions, while cytosolic marker EF1α largely remained largely in the supernatant (**Fig. 6C**). Immunofluorescence analysis of 1×TY::ESAG3^RR^ showed extensive colocalisation with BiP (**Fig. 6D, top**), and live-cell imaging of C-terminally tagged ESAG3::mNeonGreen revealed a reticular ER pattern (**Fig. 6D, bottom**) with no plasma membrane signal. ESAG3 lacks predicted transmembrane domains or GPI-anchor signals, consistent with a soluble ER-luminal glycoprotein. Together, these data establish ESAG3 as an *N-*glycosylated ER-resident glycosyltransferase, positioned to encounter VSG substrates during ER transit and maturation.

### Recognition of *O*-glycan-dependent Ab250 epitope activates complement-mediated trypanolysis

Having established ESAG3 as the VSG3 *O-*glucosyltransferase, we asked whether the *O*-glucosylation-dependent Ab250 epitopes could activate the classical complement pathway and lyse bloodstream-form trypanosomes. To exclude Ig-mediated cytotoxicity at high antibody titres, we determined that 50 µg/mL was the maximum concentration that did not cause complement-independent lysis (**Fig. S9**). Next, we examined the distribution of fluorescently labelled Ab250 on VSG3-WT cells at steady state: at 37°C in the absence of complement, Ab250 signal was largely absent from the cell surface, consistent with hydrodynamic flow-mediated clearance^47,48^; in the presence of complement, fluorescence accumulated as an intense persistent posterior focus (**Fig. 7A, B**), indicating that complement likely alters the fate of Ab250-VSG3 immune complexes at the trypanosome surface.

**Figure 7.**
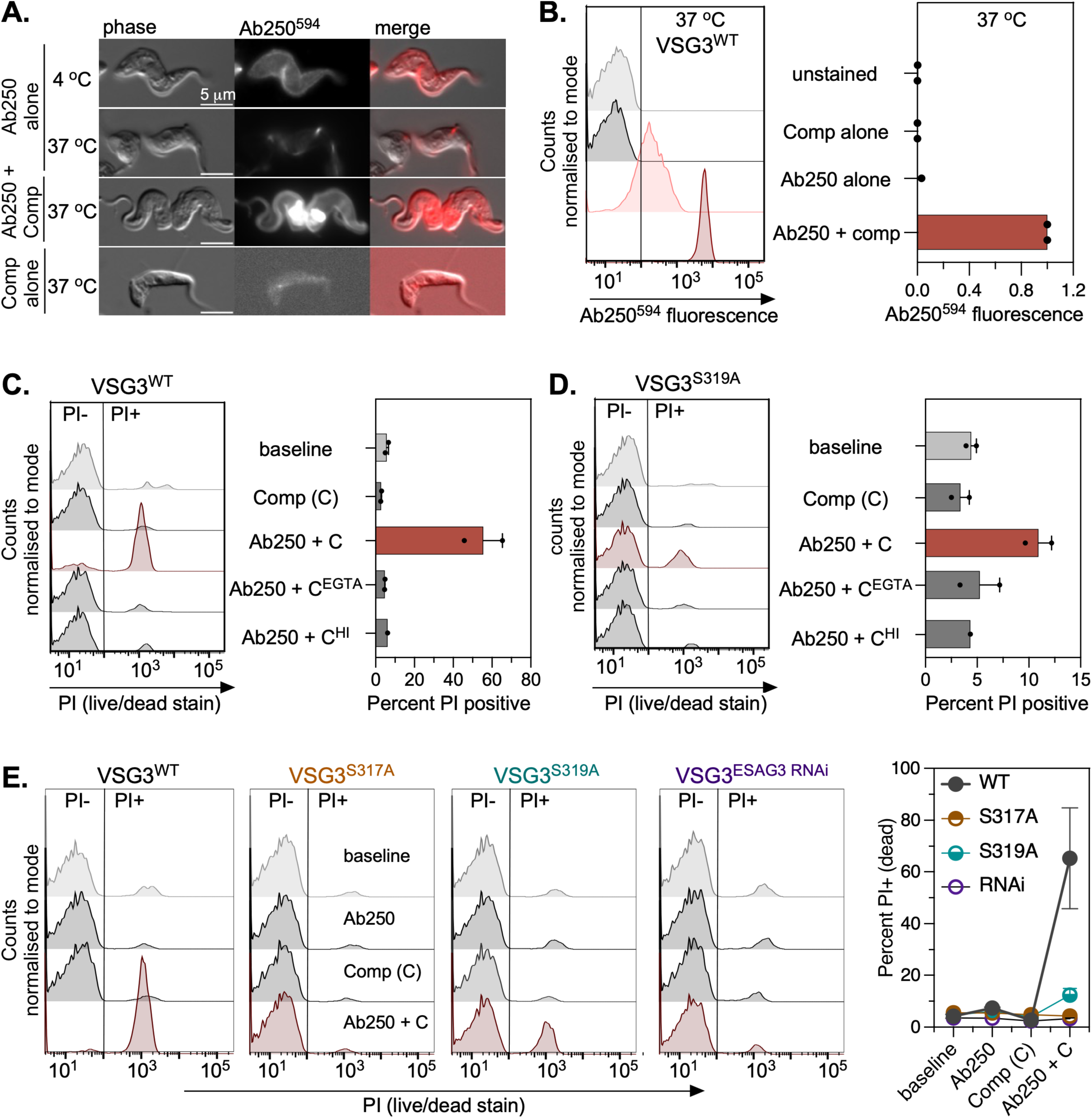
ESAG3-dependent *O*-glucosylation of VSG3 determines susceptibility. **A.** Live-cell fluorescence microscopy of VSG3^WT^ trypanosomes labelled with Ab250-Alexa Fluor 594 (Ab250⁵⁹⁴, red). Cells were incubated with Ab250^594^ at 4°C (Ab250 alone, 4°C) or shifted to 37°C in the presence or absence of guinea pig complement (Ab250 + comp; comp alone) for 25 min, then fixed immediately. Phase contrast, Ab250^594^ fluorescence, and merged images are shown. Scale bar = 5 µm. Representative of two independent experiments. **B.** Quantification of Ab250^594^ surface fluorescence intensity (MFI) under the indicated conditions. Ab250-VSG3 immune complexes are retained at the cell surface when complement is present relative to Ab250 alone, consistent with complement-mediated impediment of surface clearance. Individual data points and means shown; n = 2 independent experiments, 10,000 single events per experiment. **C, D.** Complement-mediated trypanolysis of VSG3^WT^ **(C)** and VSG3^S319A^ **(D)** cells requires active complement. Bloodstream-form trypanosomes were incubated without Ab250 antibody or complement (baseline); complement alone (C); Ab250 + complement (Ab250 + C); Ab250 + EGTA/EDTA-inactivated complement (Ab250 + C^EGTA^); Ab250 + heat-inactivated complement (Ab250 + C^HI^, 56°C, 30 min). Cell viability was assessed by propidium iodide (PI) exclusion by flow cytometry, gated on single trypanosomes. Representative PI fluorescence histograms (left) and quantification of % PI-positive cells (right) are shown. Bars represent means; individual data points from two independent, 10,000 single events per experiment. **E.** Ab250-mediated trypanolysis correlates with *O-*glucose occupancy of VSG3. Trypanosomes expressing VSG3^WT^, VSG3^S317A^, VSG3^S319A^, or VSG3 ESAG3 RNAi (VSG3^ESAG3 RNAi^) were incubated with Ab250 alone; complement alone; Ab250 + complement. Assay conditions as in (C). Representative PI fluorescence histograms (n = 10,000 total events/sample) for each cell line (left); quantification of % PI-positive cells (right); n = 2 independent experiments. Note: We observed loss of lysed cells from the single-cell event gates in VSG3^WT^ Ab250 + complement samples (n = ∼5,555–6,809 events in WT vs ∼89,141–9,217 in controls); suggesting the true extent of complement-mediated lysis is underestimated and may approach 60–90% in VSG3^WT^ cells.

To determine whether Ab250-opsonised trypanosomes could activate the classical complement pathway, we incubated VSG3-WT and VSG3-S319A cells with Ab250 and guinea pig complement. VSG3-WT and VSG3-S319A cells retain 100% and ∼32% of wild-type Ab250 surface binding respectively (**Fig. 1D**). VSG3-WT cells were efficiently lysed (mean 55.6% PI-positive) while VSG3-S319A cells, retaining partial *O-*glycan occupancy at S317, showed markedly reduced lysis (mean 10.9% PI-positive); neither Ab250 alone nor complement alone increased cell death above the 3–7% baseline (**Fig. 7C, D**). Lysis was abolished by EGTA/EDTA chelation and heat inactivation of complement, confirming dependence on active Ca^2+^-dependent complement components and establishing that lytic efficiency correlates with Ab250 surface binding (**Fig. 7C, D**).

To determine whether trypanolysis correlates with *O-*glycan occupancy, we tested all four cell lines spanning the full range of Ab250 surface binding (**Fig. 1D,F**). Cells with relatively full Ab250 binding (VSG3-WT) were lysed at high efficiency; cells with partial binding (VSG3-S319A, ∼32% of wild-type) showed proportionally reduced lysis; and cells lacking S317 *O-*linked glycan entirely — whether through point mutation (VSG3-S317A) or ESAG3 depletion — were not killed above background (**Fig. 7E**). The identical protection of VSG3-S317A and ESAG3 RNAi cells confirms that complement resistance is conferred specifically by the absence of ESAG3-catalysed *O-*glycosylation-dependent Ab250 epitope at S317, not a structural effect of the serine-to-alanine substitution.

## Discussion

Antigenic variation in African trypanosomes is understood classically as a process in which VSG switching generates sequence polymorphisms to diversify antibody epitopes for immune evasion. Previous work expanded this view, showing that heterogeneous *O-*glycosylation of hypervariable surface-exposed loops on VSG3 reshapes antibody repertoires and influences host survival^20,23^. Yet the enzyme generating this immunologically relevant modification remained elusive. Here, we identify ESAG3 as the *O*-glucosyltransferase required for generating these diverse antibody epitopes on VSG3. To our knowledge, ESAG3 is the first protein *O*-glucosyltransferase known to generate an α-*O*-glucose-serine linkage on a eukaryotic surface antigen. The co-expression of ESAG3 with VSGs within the same polycistronic transcription unit couples *O*-glucosylation with monoallelic allelic VSG expression, integrating two strategies of antigenic variation into a unified system.

Beyond its role in generating glycoform heterogeneity, ESAG3-dependent *O-*glucosylation has opposing consequences at the host-trypanosome immune interface. We show that the same modification that limits antibodies focusing on immunodominant VSG loops^23^, can also create a glycan-dependent antibody epitope whose recognition triggers complement-mediated trypanolysis (**Fig. 7**). Importantly, VSG3-WT cells are efficiently lysed; VSG3-S319A cells, retaining approximately 32% of WT Ab250 binding, are less susceptible; and VSG3-S317A and ESAG3-depleted cells, with Ab250 binding reduced to background, are not killed above background. Recognition of the glycan-dependent epitope is therefore a determinant of complement susceptibility under these experimental conditions. The requirements for Ab250 and Ca^2+^ are consistent with activation of the classical complement pathway. Lysis occurs despite hydrodynamic sorting of antibody–VSG complexes towards the flagellar pocket^47,48^. In the presence of complement, Ab250–VSG3 complexes persist at the posterior pole, suggesting that complement engagement delays their removal sufficiently for lysis to proceed. These effects are biologically opposing but mechanistically compatible: population-level glycoform heterogeneity limits antibody-repertoire focusing^23^, while recognition of a particular glycan-dependent antibody epitope renders parasites displaying it susceptible to immune-effector killing. Our experiments use a single recombinant monoclonal antibody and guinea-pig complement, and do not establish whether this activity generalises to the polyclonal responses generated during experimental murine infection^23^. Whether complement susceptibility is an inadvertent cost of glycosylation-dependent diversification, or contributes to clearance of recognised variants, is an open question raised by our findings and the subject of ongoing investigations in our laboratory.

Our structural and biochemical analyses reveal an unusual relationship between ESAG3 quaternary organisation and glycosyltransferase activity. ESAG3 adopts a canonical GT-A catalytic fold, is specific for UDP-glucose and Mn^2+^-dependent, and generates an α-O-glucose linkage, identifying it as a retaining protein *O*-glucosyltransferase. Its quaternary organisation, however, is unprecedented among characterised glycosyltransferases: 18 protomers assemble into a C3-symmetric ring that organises complete catalytic units within an extensive network of inter-subunit contacts. Large ESAG3 complexes are detected both in parasite extracts and following heterologous expression, indicating that higher-order assembly is an intrinsic property of the enzyme. In many glycosyltransferases, dimeric or tetrameric assemblies are the predominant functional states and oligomerisation is frequently required for full enzymatic activity^49^.

ESAG3 extends this principle into a substantially higher-order architecture. Perturbation of the H54/W345 inter-subunit (dimer) interface disrupts this assembly and generates lower-order oligomeric species with markedly different activities, including one with substantially increased substrate conversion. LC-MS analysis further shows that H54A/W345A alters the glucosylated product distribution, with an increased relative contribution from diglucosylated peptide. *In vivo*, ESAG3 depletion abolishes VSG3 *O*-glycosylation and complementation restores heterogeneous glycoform distributions, establishing a physiological role for ESAG3 in generating VSG glycan heterogeneity. Together, our findings identify ESAG3 as a retaining GT-A *O*-glucosyltransferase with an unusual quaternary organisation. Whether this architecture controls the extent or distribution of VSG *O*-glucosylation *in vivo* remains to be determined.

Although the mechanism for generating higher-order VSG glycoforms (0/1/2/3) at S317 remains unresolved, our data demonstrate that ESAG3 initiates glucose addition on native VSG3 *in vivo*, but the enzyme(s) responsible for sugar chain extension (up to 3-hexoses in VSG3 and 2-hexoses in VSG615) is/are unknown. The *T. brucei* genome encodes a large family of 50 intact ESAG3-related (GRESAG3) genes distributed across subtelomeric and internal chromosomal regions^35,38^, raising the possibility of a multi-enzyme relay mechanism. In this model, ESAG3 would prime VSG substrates at S317, and GRESAG3 enzymes would extend the monosaccharide. Sequential extension has precedent in the Notch *O*-glucose pathway, in which POGLUT1 initiates β-*O-*glucosylation of EGF repeats and distinct downstream glycosyltransferases sequentially extend the modification^50,51^. Resolving these mechanisms will require site- and linkage-resolved mass spectrometry and biochemical testing of GRESAG3 enzymes against unmodified and mono-glucosylated VSG substrates.

Unlike core *N*-glycosylation enzymes which modify almost all VSGs^14,18,46^, ESAG3 genes are located within expression sites and are therefore subject to monoallelic expression site control^52^. Why an *O*-glycosyltransferase is encoded within the same transcriptional unit as the VSG remains an open question. Intact ESAG3 genes occur in nine of the fifteen Lister 427 bloodstream expression sites. During transcriptional switching, activation of a new expression site can change both the VSG and the ESAG3 variant expressed, whereas gene conversion can replace the VSG while the same expression site and ESAG3 gene remain active. Whether ESAG3 modifies other Class B VSGs beyond VSG3, and whether ESAG3 variants in different BESs differ in substrate specificity, remain to be established. If ESAG3 variants have distinct specificities, transcriptional switching could change both the VSG expressed and the heterogeneity of its *O*-glycans, further diversifying the epitopes presented to host antibodies.

## Acknowledgements

We thank Samuel Dean (University of Warwick, UK) for sharing the pDRv0.5 RNAi plasmid. We are grateful to Nina Papavasiliou and Erec Stebbins (DKFZ, Heidelberg) for sharing VSG3-expressing wild-type, S317A, S319A, and SSAA mutant cell lines and Ab250 monoclonal antibody, and for valuable discussions. We acknowledge Diamond Light Source for time on Titan Krios m07 at the electron Bio-Imaging Centre (eBIC) under proposal bi33230-16. Some mass spectrometry analysis was performed at the University of York. The York Centre of Excellence in Mass Spectrometry was created thanks to a major capital investment through Science City York, supported by Yorkshire Forward, the Northern Way Initiative, and EPSRC (EP/K039660/1; EP/M028127/1). We thank Aurélien F. A. Moumbock (University of Freiburg, Germany) for support with flexible docking. We thank Jay Bangs (University at Buffalo, USA). Mark Carrington (University of Cambridge) and Matt Higgins (University of Oxford, UK) for helpful comments on the manuscript. We are particularly grateful to Keith Gull (University of Oxford, UK and Imperial College London, UK) for his sustained intellectual engagement with this project, which shaped key aspects of the interpretations and conclusions of this work.

This research was funded by a Wellcome Trust and Royal Society Sir Henry Dale Fellowship (208780/Z/17/Z) to C.T. and UKRI Future Leaders Fellowship (MR/Y01975X/1) to S.L.R.

## Copyright statement

This research was funded in whole, or in part by the Wellcome Trust [Grant numbers 208780/Z/17/Z]. For the purpose of open access, the author has applied a CC BY public copyright licence to any Author Accepted Manuscript version arising from this submission.

## Author contributions

This project was conceptualised by EH and CT. QZ, JDB, CGC, LM, OWP, LG, WS, EN, AG, and CT conducted experimental work. QZ, LG, and CT performed antibody binding and complement assays. QZ purified recombinant proteins and performed biochemical assays. JDB and QZ prepared cryo-EM grids and collected cryo-EM data. QZ processed the cryo-EM data, performed 3D reconstruction, model building and validation, with advice from JDB. QZ, CT, MAJF and EH analysed glycosyltransferase assay and quantitative mass spectrometry data. AG, EP and SAV purified Ab250 antibody. MAJF, SRL, EH and CT provided resources. Project planning and supervision were done by SRL, MAJF, EH and CT. Data visualisation was prepared by QZ, MAJF, EH, and CT. CT wrote the original draft with input from QZ, MAJF, and EH. All authors approved the final manuscript.

## Methods

### Parasite culture and cell lines

Bloodstream-form *Trypanosoma brucei brucei* Lister 427 MITat1.2 (VSG221) single-marker (SM) cells were used for all experiments^53^, except for experiments requiring VSG3 (MITat1.3; VSG224) – expressing cells described below. Cells were cultured in HMI-9 medium supplemented with 10% (v/v) heat-inactivated fetal bovine serum (Gibco), 100 U mL⁻¹ penicillin, and 100 μg mL⁻¹ streptomycin (Gibco) at 37°C; at 5% CO₂. Cell density was maintained between 1 × 10⁵ and 2 × 10⁶ cells mL⁻¹. Parental SM cells were maintained under selection with G418 (2.5 μg mL⁻¹). Transgenic cell lines carrying inducible RNAi constructs or epitope-tagged ESAG3 were selected with hygromycin B (5.0 μg mL⁻¹) and/or puromycin (0.2 μg mL⁻¹) as appropriate. *T. brucei* expressing VSG3, VSG3-S317A, VSG3-S319A, and VSG3-SSAA have been described previously^20,23^.

### Construction of plasmids

The ESAG3 coding sequence (gene ID: Tb427.BES40.10) was amplified from *T. brucei* Lister 427 genomic DNA by PCR using Phusion High-Fidelity DNA Polymerase (New England Biolabs). For C-terminal fluorescent tagging, the ESAG3 open reading frame (excluding the stop codon) was tagged using CRISPR-Cas9 with pRExT2A-CT-PTy (Addgene ID: 232203) ^54^ to generate ESAG3::mNeonGreen::T2A::PAC. The endogenous 3ʹ UTR was preserved.

For RNAi knockdown, an 864 bp fragment (nucleotides 221-1085) of the ESAG3 open reading frame was cloned into the doxycycline-inducible RNAi vector pDRv0.5 and p2T7-177 stem-loop RNAi vector^55,56^. pDRv0.5 generates two copies of the target fragment in reverse complement separated by a 150-nucleotide stuffer sequence. Opposing T7 promoters under doxycycline control drive transcription of the resulting stem-loop dsRNA. The construct was linearized with NotI and targeted to the ribosomal RNA spacer array under hygromycin B selection. For RNAi experiments, knockdown was induced by addition of doxycycline (1 μg mL⁻¹; Sigma-Aldrich) to the culture medium. Growth curves were determined by daily cell counting using a hemocytometer.

For RNAi-resistant complementation, the ESAG3 coding sequence was recoded to introduce silent mutations at wobble positions throughout the targeted RNAi region while maintaining optimal codon usage for trypanosomes. The recoded sequence was synthesised (Genewiz, UK) with an N-terminal Ty1 epitope tag and cloned into pXS6 expression vector^57^. The RNAi-resistant construct (1×TY::ESAG3^RR^) was linearized with NotI, targeted to the ribosomal spacer region, and selected with puromycin.

### Trypanosome transfection and selection

Bloodstream form trypanosomes were transfected by electroporation^58^. Briefly, log phase cells (1 × 10^6^ cells mL^-1^) were harvested by centrifugation (800 × g, 10 min, room temperature), and resuspended in 1x Roditi buffer containing 10 μg linearized plasmid DNA. Cells were electroporated using program X-001 of the Amaxa Nucleofector IIb (Lonza) electroporator. Transfected cells were transferred to 10 mL HMI-9 medium and allowed to recover for 6 – 18 h overnight before addition of the appropriate selection drug. Stable cell lines were obtained after 7-10 days of selection.

### Recombinant protein expression and purification

The mature ESAG3 coding sequence (residues 23-368, lacking the native N-terminal signal peptide predicted by SignalP v5.0) was codon-optimised for mammalian expression, modified with an N-terminal CD33 signal peptide for secretion, and synthesised with a C-terminal TEV protease cleavage site, EPEA and 10×His tag (GenScript Biotech, UK). The gene was cloned into pcDNA3.1(+) vector by GenScript and site-directed mutants (D110N, W242A, H54A/W345A) were generated and sequence-verified.

Expi293F cells (ThermoFisher, A14635) were cultured in FreeStyle 293 expression medium (ThermoFisher) at 37°C in a humidified atmosphere containing 8% CO₂ in 125 mL Corning Erlenmeyer flasks with orbital shaking at 125 rpm in a New Brunswick S41i rotating incubator (Eppendorf). Cells were maintained between 0.5 × 10⁶ and 2.5 × 10⁶ cells mL⁻¹.

For transfection, plasmid DNA (200 μg) and polyethylenimine (PEI) MAX (40 kDa (Polysciences), 600 μL of 1 mg mL⁻¹ stock) were each diluted separately in Opti-MEM with GlutaMAX (ThermoFisher) to a total volume of 10 mL and incubated for 5 min at room temperature. The two solutions were combined, mixed gently, and incubated for 30 min at room temperature to allow complex formation. This transfection mixture was added dropwise to 200 mL of Expi293F cells at a density of approximately 2 × 10⁶ cells mL⁻¹. Cultures were incubated for 5 days at 37°C, 125 rpm, and 8% CO₂ for protein expression. Culture supernatants were harvested by centrifugation (200 × g, 10 min) to pellet cells, followed by clarification at 4,000 × g for 20 min at 4°C, and filtration through 0.45 μm polyethersulfone filters (Sartorius).

Recombinant ESAG3 was purified from clarified supernatants by immobilized metal affinity chromatography using a 1 mL HisTrap^TM^ Excel column (Cytiva) on an ÄKTA Pure FPLC system at 4°C. Supernatants were supplemented with 1 M HEPES pH 7.5 to a final concentration of 25 mM and 2 M imidazole pH 7.5 to a final concentration of 10 mM. All buffers were degassed and filtered through 0.22 μm filters (Sartorius). The column was equilibrated in running buffer (150 mM NaCl, 25 mM HEPES pH 7.5) and supernatants were loaded at 0.5 mL min⁻¹. The column was washed sequentially with 15 mL each of 4%, 6%, and 8% elution buffer (500 mM imidazole, 150 mM NaCl, 25 mM HEPES pH 7.5) to remove weakly bound contaminants. Protein was eluted with 100% elution buffer and concentrated to 500 μL using Vivaspin 20 centrifugal concentrators (10 kDa MWCO; Sartorius).

Concentrated eluate was further purified by size-exclusion chromatography using a Superdex 200 Increase 10/300 GL column (Cytiva) equilibrated in running buffer (150 mM NaCl, 25 mM HEPES pH 7.5) on an ÄKTA Pure system at 4°C. The column was calibrated using Gel Filtration Calibration Kit HMW (Cytiva). Samples (500 μL) were injected and eluted at 0.8 mL min⁻¹. Peak fractions corresponding to the major ∼700 kDa oligomer were pooled and concentrated to 200 μL using Vivaspin 6 centrifugal concentrators (10 kDa MWCO; Sartorius). Protein concentration was determined by NanoDrop (Thermo Fisher, UK). Samples were either used immediately for cryo-EM grid preparation or supplemented with glycerol to a final concentration of 10% (v/v), flash-frozen in liquid nitrogen, and stored at −80°C. Protein purity was assessed by SDS-PAGE with Coomassie Brilliant Blue R-250 staining, and identity was confirmed by anti-ESAG3 Western blotting.

### Blue-native PAGE

Blue-native polyacrylamide gel electrophoresis (BN-PAGE) was performed using the NativePAGE Bis-Tris Gel System (ThermoFisher). Bloodstream form trypanosomes (1 × 10⁸ cells) were harvested by centrifugation (800 × g, 10 min, 4°C), washed twice in HBSG (20 mM HEPES pH 7.4, 150 mM NaCl, 10 mM glucose), and solubilised in NativePAGE Sample Buffer (ThermoFisher) supplemented with 10% (v/v) glycerol, 1% (w/v) n-dodecyl-β-D-maltoside (DDM), 1× cOmplete protease inhibitor cocktail (Roche), and 100 μg mL⁻¹ DNase I (Roche). Samples were incubated on ice for 30 min, then centrifuged (13,000 × g, 1 h, 4°C) to remove insoluble material. Supernatants were either used directly or treated with 4 M urea to denature protein complexes. Samples (15 μL) were mixed with NativePAGE Sample Buffer containing Coomassie G-250 dye at a 4:1 sample-to-dye ratio and loaded onto precast 4-16% NativePAGE Bis-Tris gradient gels (ThermoFisher). Electrophoresis was performed at 4°C in NativePAGE Running Buffer according to the manufacturer’s instructions. After electrophoresis, proteins were transferred to PVDF membranes (Merck Millipore) by BioRad Transblot and analysed by western blotting with anti-ESAG3 antibodies. NativeMark Unstained Protein Standard (ThermoFisher) was used for molecular weight estimation.

### Cell fractionation

Bloodstream form trypanosomes (1 × 10^7^ cells) were harvested by centrifugation (800 × g, 10 min, 4°C), washed in ice-cold trypanosome dilution buffer HBSG as described in Tiengwe et al 2017^16^, and hypotonically lysed in distilled H₂O supplemented with cOmplete protease inhibitor cocktail (Roche) for 10 min on ice. Lysates were centrifuged at 13,000 × g for 10 min at 4°C to separate cytosolic (supernatant) and organelle-associated/membrane (pellet) fractions. An aliquot of the lysate prior to centrifugation was retained as the total fraction. Equal cell equivalents (1 × 10⁷ cells per lane) of total, cytosolic, and organelle-associated fractions were resuspended in 1× LDS Sample Buffer (ThermoFisher) containing 50 mM DTT, heated at 95°C for 5 min, and analysed by SDS-PAGE and Western blotting with appropriate antibodies.

### Peptide substrates

Synthetic peptides corresponding to *O-*glycosylation sites in trypanosome Class B VSGs were purchased from QYAOBIO (ChinaPeptides). All peptides were C-terminally amidated (-NH₂). VSG3 peptides contained the cysteine-flanked loop region (CTGSASEGLC, residues 314-323) with naturally occurring serine residues at positions 317 and 319. Wild-type VSG3 peptide (DD-ATGCTGSASEGLCVEY-KK-NH₂) contains charge-complementary flanking sequences (DD…KK) to enhance hydrophilicity. The VSG3 S317A mutant (DD-ATGCTGAASEGLCVEY-KK-NH₂) has serine 317 replaced with alanine to assess site-specific glycosylation. We also synthesised VSG3-2.0 peptide (KGCTGSASEGLCGGYK-NH₂) containing the same 314-CTGSASEGLC-323 loop sequence but with altered flanking regions lacking charge complementarity. Additional VSG-derived peptides included the metacyclic VSG VSG1954 (DD-ATGCDGIGGSNGGKC-KK-NH₂) and VSG615 (DD-ATGCNGGTSSAAC-KK-NH₂). SetA consensus motif peptides, SetA-SGLP (DGGSGLPGGYK-NH₂) and SetA-TGLP (DGGTGLPGGYK-NH₂), contained serine and threonine O-glycosylation motifs, respectively. For all cysteine-containing peptides, variants with and without disulfide bonds were tested to assess the importance of disulphide loop structure for ESAG3 substrate recognition and enzymatic activity.

### UDP-Glo^TM^ glycosyltransferase assay

A UDP-Glo™ Glycosyltransferase Assay kit (Promega) was used for GT activity of wild-type and mutant ESAG3 according to the supplier’s protocol. ESAG3 variants tested included wild-type enzyme, D110N, W242A and the H54A/W345A double mutant, which affects the ESAG3 oligomerization interface. Reactions were assembled in 96-well white plates (Greiner) at room temperature with a final volume of 25 µL per well. A UDP standard curve was established using a twofold dilution series of UDP from 25 µM to 0 µM. Each 25 µL reaction mixture consisted of 2.5–4.73 µM purified ESAG3, 125 µM peptide acceptor substrates, and 200 µM UDP-glucose (Promega), prepared in reaction buffer containing 25 mM HEPES (pH 7.5), 150 mM NaCl, and 5 mM MnCl₂. Reactions were incubated at 30 °C for 1 hr or overnight at RT, followed by the addition of 25 µL UDP-Glo^TM^ nucleotide detection reagent. After a further 1-hour incubation at room temperature, luminescence signals were measured using a Tecan SPARK multimode microplate reader. The range of measurements was determined to be in the linear range of detection, where luminescence is directly proportional to UDP concentration.

### High-Performance Liquid Chromatography (HPLC) assay

Glycosyltransferase activity of recombinant wild-type and mutant ESAG3 proteins was detected using a C18 reversed-phase HPLC column (Poroshell 120 EC-C18, Agilent) on an Agilent 1260 Infinity II system with Chemstation software. The column was equilibrated in 90% buffer A (Chromplete water with 0.1% trifluoroacetic acid, TFA) and 10% buffer B (acetonitrile with 0.085% TFA) at a flow rate of 0.5 mL/min. Each reaction mixture contained 5 µM purified ESAG3, 125 µM peptide substrates, and 200 µM UDP-glucose in glycosyltransferase buffer (pH 7.5, 25 mM HEPES, 150 mM NaCl, 5 mM MnCl₂). After running a blank sample, reaction mixtures were injected directly onto the column. Peptide separation was performed using a gradient of buffer B from 10% to 30% over 22 minutes at a flow rate of 0.5 mL/min, followed by column re-equilibration. Detection was performed by UV absorbance at 280 nm. For each peptide variant, modified and unmodified peptide species were quantified using peptide-specific elution windows. Peak areas were determined from the baseline-corrected area under the UV280 absorbance–time curve (AUC), using consistent integration parameters across samples within each peptide series. Percentage conversion was calculated as: Conversion (%) = ΣAUC (product peaks) / [Σ AUC (product peaks) + AUC (unmodified substrate)] × 100. Chromatograms were plotted in GraphPad Prism (v11). For traces with flat baseline in the expected product elution window, conversion was recorded as 0%.

### Mass spectrometry

MALDI–TOF mass spectrometry was performed on HPLC fractions at the University of York Metabolomics & Proteomics Facility. For each sample, 1 μL of material was mixed 1:1 with α-cyano-4-hydroxycinnamic acid (10 μg μL⁻¹), spotted onto a MALDI target plate, and air-dried. Spectra were acquired over a mass range of m/z 800–4000 in reflection mode, with matrix suppression active below m/z 650.

For LC–MS/MS analysis, samples were loaded onto EvoTip Pure tips (Evosep) and separated on an 8 cm Performance C18 column using a 300 samples-per-day (SPD) preset gradient on an EvoSep UPLC system. Data-dependent acquisition parallel accumulation–serial fragmentation (DDA-PASEF) was performed on a Bruker timsTOF HT mass spectrometer. Blank runs were included between each sample to minimise carryover.

Raw LC-MS/MS data were searched using FragPipe (v23.1) against a custom database containing VSG3, VSG3-mutant, VSG615 and VSG1954 sequences, together with common proteomic contaminants. Searches were performed at a 1% false discovery rate (FDR). Hexose modifications (+162.05 Da) on serine and threonine residues were included as variable modifications. Modification site localisation probabilities were assigned by FragPipe. Peptide spectral matches (PSMs) and relative signal intensities were extracted from FragPipe output and used as measures of MS/MS evidence for peptide modification. High-confidence modifications were defined as those supported by ≥2 PSMs with unambiguous site assignment (localisation score = 1.0).

### VSG3 purification, in gel digestion and LC-MS analysis of glycopeptides

WT and ESAG3 RNAi VSG3-expressing cells were washed with ice cold HEPES-buffered saline containing glucose (HBS; 50 mM HEPES/KOH [pH 7.5], 5 mM KCl, 50 mM NaCl, 20 mM glucose). VSG3 was purified as described in^1,59^. Samples of purified VSG3 (4 mg) were dissolved in 10 ul SDS-sample buffer, adjusted to 10 mM dithiothreitol and reduced for 10 min at 70°C, adjusted to 50 mM iodoacetamide and alkylated for 30 min at room temperature in the dark. Samples were subjected to SDS-PAGE on 4-12% pre-cast gels (Invitrogen), stained with Generon QC stain (Invitrogen) and the prominent VSG bands at around 60 kDa were excised transferred to microcentrifuge tubes and cut into small pieces. Gel pieces were washed sequentially for 15 min each with 200 μL of Milli-Q water, 200 μL of acetonitrile, 200 μL of 100 mM ammonium bicarbonate, and 200 μL of 100 mM ammonium bicarbonate/acetonitrile (50:50, v/v). Finally, the gel pieces were washed with 100 μL of acetonitrile for 10 min and briefly dried in a SpeedVac. In-gel digestion was carried out with trypsin at a final concentration of 12.5 μg/mL in 20 mM ammonium bicarbonate, followed by incubation at 30 °C for 16 h with gentle shaking. An equal volume of acetonitrile was added, and peptides were extracted by shaking at 30 °C for 15 min. The resulting supernatant was transferred to a fresh microcentrifuge tube. Residual peptides were further extracted from the gel pieces by adding 5% formic acid and shaking for 15 min, followed by the addition of an equal volume of 100% acetonitrile and shaking for 45 min. The supernatants were pooled, and the gel pieces were washed once more with acetonitrile for 10 min. Supernatants from each step were combined, dried in a SpeedVac, and stored at –20 °C.

VSG3 tryptic peptides were analysed using Exploris 480 (Thermo Scientific) mass spectrometer coupled with a Vanquish nano-LC (Thermo Scientific). LC buffers were the following: buffer A and weak buffer wash (0.1% formic acid in Milli-Q water (v/v)); buffer B and strong buffer wash (80% acetonitrile, 0.1% formic acid in Milli-Q water (v/v)). Samples were loaded onto a trap column (Pep Map Neo C18, 5 µm, 300 µm x 5 mm, Thermo Scientific) equilibrated in buffer A with flow-control loading at 10 µl/min. The column (Pep Map Neo C18, 75 µm x 500 mm, Thermo Scientific) was equilibrated with 2% buffer B and kept at a constant temperature of 50°C. Peptides were eluted from the column at a constant flow rate of 300 nl/min with linear gradients from 2% buffer B to 6% buffer B in 5 min, from 6% buffer B to 35% buffer B in 65 min and finally buffer B was increased from 35%B to 96% B in 2 min. The column was then washed with 96% buffer B for 13 min. Two blanks were run between each sample to reduce carryover. Chromatography was performed using a Thermo EASY-Spray source operated in positive mode with spray voltage at 2.0 kV and the ion transfer tube temperature of 250°C. The Exploris 480 was used in data dependent mode. A scan cycle comprised MS1 scan (*m/z* range from 335 to 1800) with a maximum ion injection time of 25 ms, Orbitrap resolution 60,000, RF lens 50% and normalized AGC target 300%. The MS1 survey scan was followed by 20 data dependant MS2 scans with an isolation window set to 1.4 Da, Orbitrap resolution at 15,000, maximum ion injection time at 40 ms and HCD collision energies at 30%. The tryptic peptide representing residues 311 to 339 of the VSG3 sequence (ATGCTGSASEGLCVEYTALTAATMQNFYK) and its +1-, +2- and +3-hexose residue variants were detected by extracted ion profiling (EIP) for their second isotope [M+3H]^3+^ ions at *m/z* 1035.1332, 1089.1562, 1143.1720 and 1197.1914, respectively, with a 10 ppm window using Xcalibur software. Relative quantification of species within a sample was achieved by integration of the EIP chromatograms. EIP was also performed using *m/z* 5.332 higher values than those described above to detect and quantify [M+3H]^3+^ methionine-oxidised variants of the same (glyco)peptides.

### Antibody production and immunological reagents

Rabbit polyclonal anti-ESAG3 antibodies were raised against the synthetic peptide C-DAVKENDLFKAKKL corresponding to residues 355-368 of full length ESAG3 (GenScript, UK). Additional antibodies described in our previous publications^16,39^ include: mouse monoclonal anti-BiP, Rabbit polyclonal anti-HSP70, mouse monoclonal anti-Ty1 (ThermoFisher; 1:1000 for Western blot, 1:500 for immunofluorescence), anti-EF1α (ThermoFisher; 1:5000), and rabbit polyclonal anti-transferrin receptor (gift from Prof. Piet Borst, Netherlands Cancer Institute). HRP-conjugated goat anti-rabbit IgG and goat anti-mouse IgG secondary antibodies (ThermoFisher) were used at 1:10,000 for Western blotting. Alexa Fluor 488- and 594-conjugated secondary antibodies (Fisher Scientific) were used at 1:1000 for immunofluorescence. Monoclonal antibody Ab250 has been described previously^23^.

### SDS-PAGE and Western blotting

Protein samples were mixed with 2× Laemmli sample buffer (125 mM Tris-HCl pH 6.8, 4% SDS, 20% glycerol, 0.02% bromophenol blue, 100 mM DTT) and heated at 95°C for 10 min. Samples were separated on 10% SDS-PAGE gels (Bio-Rad TGX FastCast Acrylamide Kit) or precast 4-12% NuPAGE Bis-Tris gels (ThermoFisher). Electrophoresis was performed in Tris-glycine running buffer (25 mM Tris, 190 mM glycine, 0.1% SDS) at 40 V for 30 min followed by 100 V for 100 min for homecast gels and at 100-200 V for 35-45 min for precast gels. Gels were either stained with InstantBlue Coomassie Protein Stain (Bio-Rad) or transferred to PVDF membranes (Merck Millipore).

For Western blotting, proteins were transferred to 0.45 μm PVDF membranes using the Trans-Blot Turbo Transfer System (Bio-Rad) at 25 V for 10 min. Membranes were blocked in 5% (w/v) non-fat milk in PBST (PBS with 0.1% Tween-20) for 1 hour at room temperature. Primary antibodies were diluted in blocking buffer and incubated for 1 hour at room temperature or overnight at 4°C. After three 10 min washes in PBST, membranes were incubated with HRP-conjugated secondary antibodies (1:5000 for anti-ESAG3; 1:5,000 for others) for 1 hour at room temperature. After three additional PBST washes, blots were developed using SuperSignal West Pico PLUS Chemiluminescent Substrate (ThermoFisher) and imaged on a ChemiDoc Imaging System (Bio-Rad).

### Flow cytometry analysis and immunofluorescence microscopy with Ab250 monoclonal antibody

Mid-log phase bloodstream-form trypanosomes (1 × 10⁶ cells mL⁻¹) expressing VSG3-WT, VSG3-S317A, VSG3-S319A, VSG3-SSAA, or ESAG3 RNAi and rescue cell lines were harvested and chilled on ice for 1 hour. Cells were incubated with Ab250-Alexa Fluor 594 (1:500 dilution; final concentration 4 μg mL⁻¹) for 10 min at 4°C in the dark. Cells were fixed by adding an equal volume of 2% (w/v) paraformaldehyde in PBS (final concentration 1%) for 10 min at 4°C. Fixed cells were washed once in PBS, settled onto poly-L-lysine-coated coverslips for 10 min, and mounted in VectaShield without DAPI. Images were acquired on a Zeiss Axio Imager.M2 microscope with a 100× oil immersion objective (NA 1.4) using identical settings across all samples.

Flow cytometry was performed on a BD LSRFortessa analyzer using a 561 nm laser for Alexa594 excitation and 610/20 nm bandpass filter for emission. Forward scatter (FSC) and side scatter (SSC) parameters were used to gate single cells. A minimum of 10,000 single-cell events per sample were acquired. Data were analysed using FlowJo v10.8 software. Median fluorescence intensity (MFI) was calculated for each sample and normalized to VSG3-WT cells. Background fluorescence was determined using parental VSG2-expressing cells incubated with Ab250-Alexa594 under identical conditions. Data represent mean ± SD from three to six independent biological experiments.

### Cryo-electron microscopy sample preparation

Purified recombinant ESAG3 oligomers (0.64 mg mL⁻¹) from the high-molecular-weight SEC peak were used for cryo-EM grid preparation. Quantifoil R2/2 300-mesh holey copper grids (Quantifoil Micro Tools) were glow-discharged for 90 s using a plasma cleaner. Sample (3 μL) was applied to grids using a Vitrobot Mark IV (ThermoFisher) operated at 4°C and 100% humidity. After a 10 s wait time, grids were blotted for 5 s and immediately plunge-frozen in liquid ethane cooled by liquid nitrogen. Vitrified grids were stored in liquid nitrogen until data collection.

### Cryo-EM data collection

Initial screening was performed on a Glacios transmission electron microscope (ThermoFisher) operated at 200 kV and equipped with a Falcon 4 direct electron detector at Imperial College London. Data were collected at 79,000× nominal magnification (1.50 Å/pixel calibrated pixel size) using EPU automated acquisition software (ThermoFisher). A total of 3,640 movies were recorded with 40 e⁻/Å² total dose fractionated into 40 frames with defocus range −1.0 to −2.5 μm.

High-resolution data collection was performed at the electron Bio-Imaging Centre (eBIC) at Diamond Light Source (UK; BAG session bi33230-16) on a Titan Krios m07 transmission electron microscope (ThermoFisher) operated at 300 kV and equipped with a K3 direct electron detector (Gatan). A total of 23,980 movies were collected at a nominal magnification of 81,000× (calibrated pixel size of 1.058 Å/pixel at the specimen level) using EPU software. Movies were recorded with 39.94 e⁻/Å² total dose fractionated into 40 frames with defocus range −0.8 to −2.5 μm. Micrographs were motion-corrected on-the-fly using RELION via the Murphy client-server architecture (Diamond Light Source) and transferred to a local server for processing.

### Cryo-EM image processing and 3D reconstruction

All image processing was performed using cryoSPARC v4.2.1. The processing pipeline is shown in **Fig. S4**. Motion-corrected micrographs (23,980) were imported and subjected to patch CTF estimation. Manual curation based on ice thickness, average defocus, CTF fit resolution, and defocus range reduced the dataset to 16,253 high-quality micrographs.

Initial particle picking was performed using Blob Picker with particle diameter 150-300 Å, generating 2,358,788 picks. These were extracted and subjected to three rounds of reference-free 2D classification to remove noise and aggregates, yielding 310,580 particles showing clear ring-shaped features. A subset of 2,368 micrographs was used for Topaz training^60^. Topaz-extracted particles underwent three additional rounds of 2D classification, resulting in 415,612 high-quality particles for 3D reconstruction.

An initial 3D reference model was generated by ab initio reconstruction in cryoSPARC using C1 symmetry. Inspection revealed clear threefold symmetry. Particles were subjected to homogeneous refinement with C3 symmetry enforced, followed by non-uniform refinement, yielding a ring structure at 3.4 Å resolution (gold-standard Fourier shell correlation at 0.143 threshold).

To improve resolution of individual six-protomer repeating units (asymmetric units of the C3-symmetric structure), C3 symmetry expansion was applied to generate three copies of each particle rotated by 120°, yielding 1,246,836 symmetry-expanded particles. A segment mask encompassing a single six-protomer unit was created in UCSF ChimeraX^61^ and applied during particle subtraction. Local refinement of the masked six-protomer unit followed by non-uniform refinement produced a final map at 3.0 Å resolution. Local resolution maps were calculated using cryoSPARC Local Resolution tool. The complete 18-protomer octadecamer comprises three six-protomer units related by C3 symmetry.

### Atomic model building and refinement

An initial homology model of the ESAG3 monomer (residues 23-368) was generated using AlphaFold3 via the AlphaFold Server. The predicted model showed high confidence (pLDDT > 90) across the structured region. For the six-protomer unit map (3.0 Å resolution), the AlphaFold monomer model was rigid-body fitted into density using UCSF ChimeraX “Fit in Map” tool. The hexameric model was refined in ISOLDE using molecular dynamics flexible fitting against the cryo-EM density, followed by manual adjustment and removal of regions lacking corresponding EM density.

The complete 18-protomer ring model was constructed by applying C3 symmetry operations to the refined hexamer to produce two additional six-protomer units. The resulting assembly was manually inspected and locally adjusted at inter-subunit interfaces in ISOLDE^62^, followed by global real-space refinement. Final optimisation of bond lengths and angles was performed using the REFMAC Servalcat function in the CCP-EM software suite v1.7.0^63^. *N*-linked glycans at position N99 were modelled using GlycoSHIELD^64^. Data collection and refinement statistics are summarised in **Table S2**.

### Antibody-mediated complement trypanolysis assay

Bloodstream-form trypanosomes were harvested by centrifugation (1,400 × g, 10 min, 4°C), washed twice in ice-cold HBSG supplemented with 0.5 µg/mL BSA, and resuspended at 1 × 10^8^ cells/mL. Cells were incubated with Ab250 (50 µg/mL) on ice for 30 min, then Guinea pig complement (Sigma-Aldrich S1639) was added at 25% final volume and cells transferred to 37°C for 30 min. For complement inactivation controls, guinea pig complement was treated with 5 mM EGTA/EDTA (final concentration) for 10 min on ice prior to addition, or heat-inactivated at 56°C for 30 min. Cell viability was assessed by propidium iodide (PI) exclusion: cells were stained with PI (1 µg/mL, 10 min at room temperature) and analysed immediately by flow cytometry as above. Events were gated on single trypanosomes (minimum 5,000 events) and PI-positive (610/20) and PI-negative populations were quantified using FlowJo v10.

To determine the minimum effective Ab250 concentration for complement-dependent trypanolysis and exclude non-specific cytotoxicity, trypanosomes were incubated with a 1:2 serial dilution series of Ab250 (200–0.5 µg/mL) in HBSG/BSA supplementation, without Guinea pig complement and assessed by PI exclusion as described above. Live-cell fluorescence microscopy of surface antibody dynamics was performed as described above.

## Data availability

Cryo-EM maps and atomic models have been deposited in the Electron Microscopy Data Bank (EMDB) and Protein Data Bank (PDB), respectively, for the ESAG3 octadecamer (EMD-59320, PDB 33BC) and the six-protomer unit obtained by focused refinement (EMD-59300, PDB 33AK). Mass spectrometry data is available in MassIVE with accession number MSV000103045 and ProteomeXchange identifier PXD083353.

