## Supporting information for "An octadecameric *O-*glucosyltransferase generates diversity in antibody epitopes on variant surface antigens in African trypanosomes"

**Competing interest statement:** The authors declare no competing interests.

Key words: *Trypanosoma brucei*, ESAG3, variant surface glycoprotein, *O*-glycosylation, antigenic variation

**This PDF includes:**

1. Figures S1–S9,
2. Tables S1–S2

**Table S1**

ESAG3 genes and pseudogenes in BES of *T. brucei* Lister 427

| | Lister 427 geneID | definition of pseudogene | VSG in BES |
|---|---|---|---|
| BES1 | Tb427.BES40.10<br>Tb427.BES40.16<br>Tb427.BES40.5pseudogene | 308 codons (3 stop codons) | VSG2 |
| BES2 | Tb427.BES129.10<br>Tb427.BES129.6pseudogene | 339 codons (6 stop codons) | VSG9 |
| BES3 | Tb427.BES4.7<br>Tb427.BES4.4pseudogene | fragment 184 amino acids | VSG6 |
| BES4 | Tb427.BES5.8<br>Tb427.BES5.4pseudogene | 255 codons (4 stop codons) | VSG21 |
| BES4** | Tb427.BES28.10<br>Tb427.BES28.6pseudogene | 339 codons (11 stop codons) | |
| BES5 | Tb427.BES98.4pseudogene<br>Tb427.BES98.8pseudogene | 339 codons (13 stop codons), 369 codons (1 stop codon) | VSG18 |
| BES7 | Tb427.BES65.8<br>Tb427.BES65.4pseudogene<br>Tb427.BES65.12pseudogene | 247 codons (17 stop codons), 307 codons (14 stop codons) | VSG3 |
| BES8 | None | None | VSG14 |
| BES10 | Tb427.BES134.5pseudogene | 245 codons (12 stop codons) | VSG15 |
| BES11 | Tb427.BES122.7<br>Tb427.BES122.4pseudogene | fragment 246 amino acids | VSG16 |
| BES12 | Tb427.BES29.5pseudogene | 232 codons (5 stop codons) | VSG8 |
| BES13 | Tb427.BES56.5pseudogene<br>Tb427.BES56.9pseudogene | fragments 244 amino acids, 291 amino acids | VSG17 |
| BES14 | Tb427.BES10.4pseudogene | fragment 212 amino acids | VSG19 |
| BES15 | Tb427.BES126.9<br>Tb427.BES126.5pseudogene<br>Tb427.BES126.14pseudogene | fragment 214 amino acids, 309 codons (8 stop codons) | VSG11 |
| BES17 | Tb427.BES51.5pseudogene<br>Tb427.BES51.8pseudogene | 239 amino acids, 368 codons (9 stop codons) | VSG13 |

Footnotes **BES4 has two Transformation-associated recombination (TAR) clones TAR5 and TAR28.
The Lister 427 geneIDs were used for this table because of better annotations than 427-2018 genome

#### Figure S1

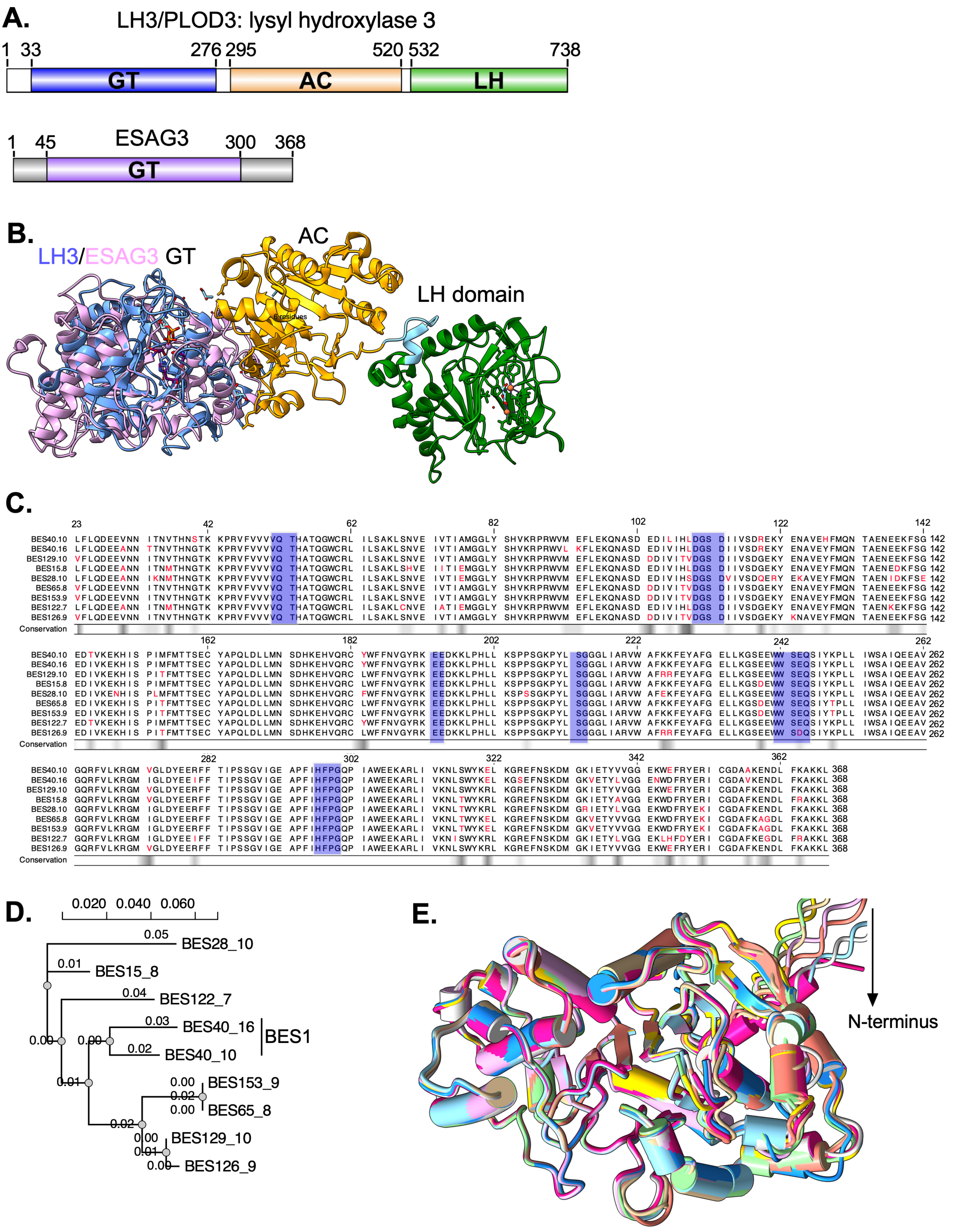


**Figure S1. ESAG3 shares structural homology with the glycosyltransferase domain of human lysyl hydroxylase 3 (LH3/PLOD3) and BES associated variants show high sequence and structural conservation**

**A. Domain architecture comparison.** LH3/PLOD3 contains three domains: glycosyltransferase (GT, blue; residues 33-276), an accessory (AC, gold; residues 276-520), and lysyl hydroxylase (LH, green; residues 520-738) domains. ESAG3 (residues 45-300 of mature protein) corresponds to the LH3/PLOD3 GT domain.

**B.** Structural superposition of AlphaFold3-predicted structure of ESAG3 (pink) with the LH3/PLOD3 GT domain (blue/purple), AC domain (gold), and LH domain (green). The ESAG3 GT-A structural core superposes with the LH3 GT domain with an RMSD of 2.40 Å over 147 Cα pairs, with no comparable structural similarity to the AC or LH domains. RMSD was calculated using PyMOL super; visualisation was generated in ChimeraX.

**C.** Multiple sequence alignment (MSA) of nine ESAG3 proteins from *T. brucei* Lister427 show conserved GT-A motifs. Conserved residues are shown in black; blue shading indicates core GT-A motifs; variable residues in red. Conservation track at bottom shows degree of amino acid conservation at each position. Alignment generated using CLC genomics Workbench v20.0.5. Predicted signal sequence (residues 1-22, SignalP v5) not shown.

**D.** Maximum likelihood phylogenetic tree of ESAG3 sequences from (**A**). Branch lengths indicate evolutionary distance (substitutions/site). BES28.10 shows the longest branch length (~0.05), while BES153.9, BES65.8, BES129.10, and BES126.9 form a closely related cluster (branch lengths ~0.00-0.01). Tree constructed from MSA using MrBayes and PhyML.

**E.** Structural superposition of AlphaFold3-predicted ESAG3 monomer structures corresponding to sequences in (**C**), aligned to BES153.9 as reference. Each monomer coloured differently. All structures align with RMSD 0.4-0.6 Å relative to BES153.9. Structures shown exclude residues 1-22 (signal sequence) and 346-368 (unstructured C-terminus). Visualisation: ChimeraX v1.10.1 Matchmaker.

### Figure S2

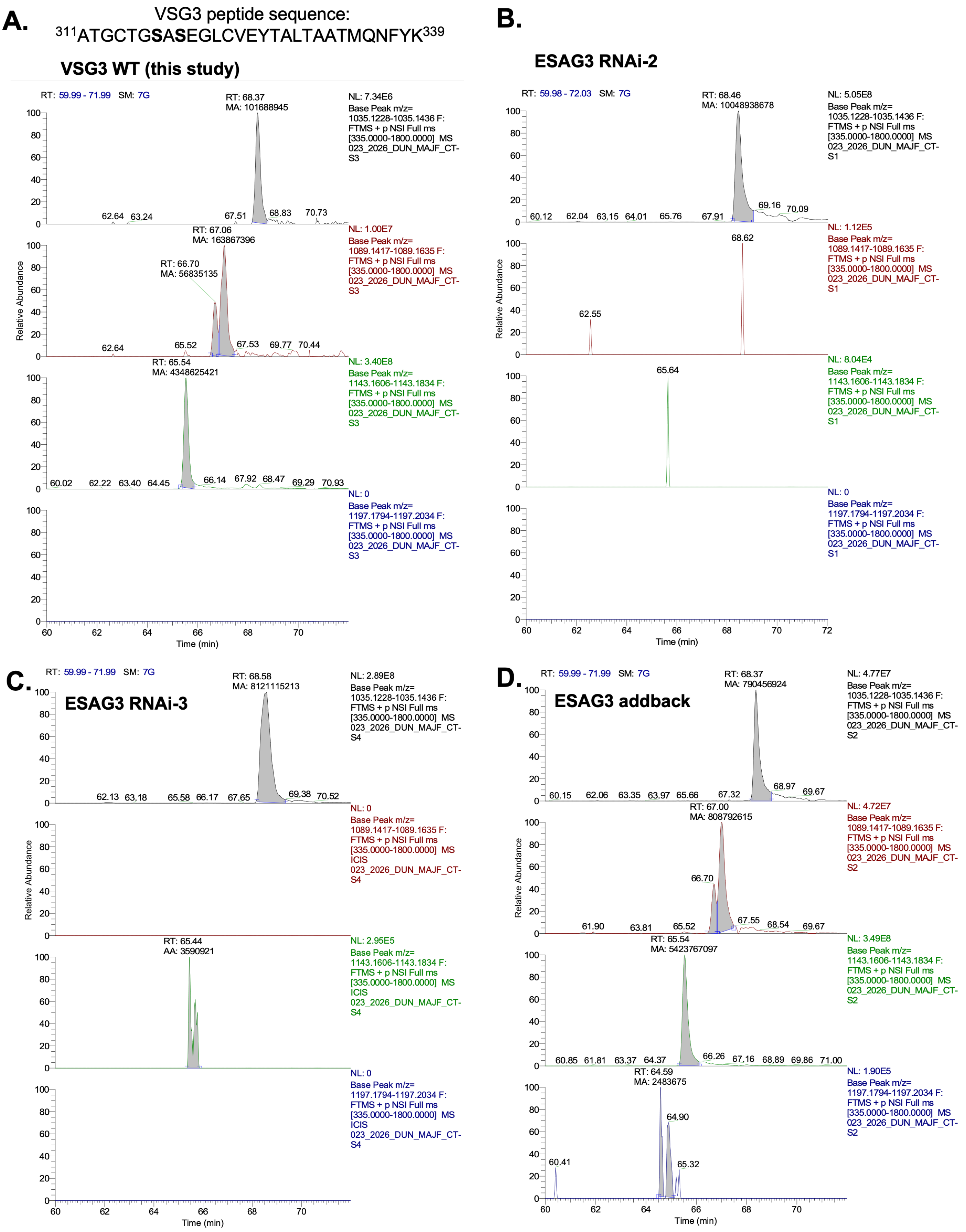


black = 0Hex, red = 1Hex, green = 2Hex, blue = 3Hex

**Figure S2. Accurate-mass extracted ion chromatograms of VSG3 glycopeptide glycoforms.**
Extracted ion chromatograms for the VSG3 tryptic glycopeptide $^{311}$ATGCTGSASEGLCVEYTALTAATMQNFYK$^{339}$ (A311–K339) from wild-type **(A)**, ESAG3 RNAi-2 **(B)**, ESAG3 RNAi-3 **(C)**, and ESAG3 addback **(D)** bloodstream-form *T. brucei* parasites. Within each panel, traces show $[M+3H]^{3+}$ extracted ion chromatograms corresponding to the unmodified peptide (0-hexose, black, m/z 1035.1228–1035.1436), and peptides carrying one (1-hexose, red, m/z 1089.1417–1089.1635), two (2-hexose, green, m/z 1143.1606–1143.1834), or three (3-hexose, blue, m/z 1197.1794–1197.2034) hexose residues. Each trace is autoscaled to maximum signal within that trace). Quantitative glycoform percentages calculated from integrated peak areas are shown in **Fig. 1F**. NL values indicate the maximum signal intensity for each trace. In wild-type parasites (**A**), the 2-hexose glycoform is dominant and the 1-hexose channel resolves two co-eluting species at RT 67.06 and 66.70 min, consistent with mono-glucosylation at S317 and S319 as distinct isoforms. ESAG3 RNAi lines (**B, C**) show near-complete loss of all glycosylated species. Addback parasites (**D**) restore the glycoform distribution, including the 1-hexose doublet, confirming specificity of the RNAi phenotype. The 3-hexose trace in the addback sample shows a low-intensity signal (NL 1.90E5; 0.03% by area). No 3-hexose signal was detected in wild-type or either RNAi line under these conditions. These experiments were performed once.

### Figure S3

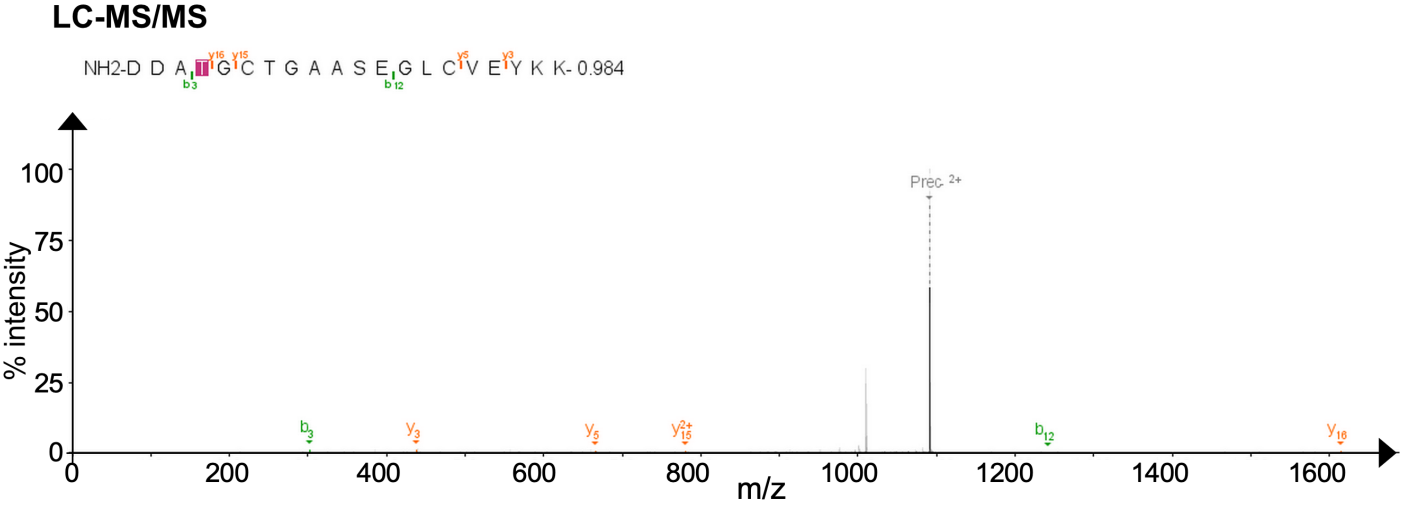


**Figure S3. LC-MS/MS site mapping of ESAG3-mediated glucosylation**
Representative LC-MS/MS spectrum showing +162.05 Da modification on VSG3 (S317A,S319) peptide at threonine 7 (T7, corresponding to T315 in full-length VSG3). Annotated fragment ions (b-ions in green, y-ions in orange) unambiguously localise glucose to T7. This modification was detected with high confidence across multiple independent peptide-spectral matches in samples containing ESAG3 and UDP-glucose.

#### Figure S4

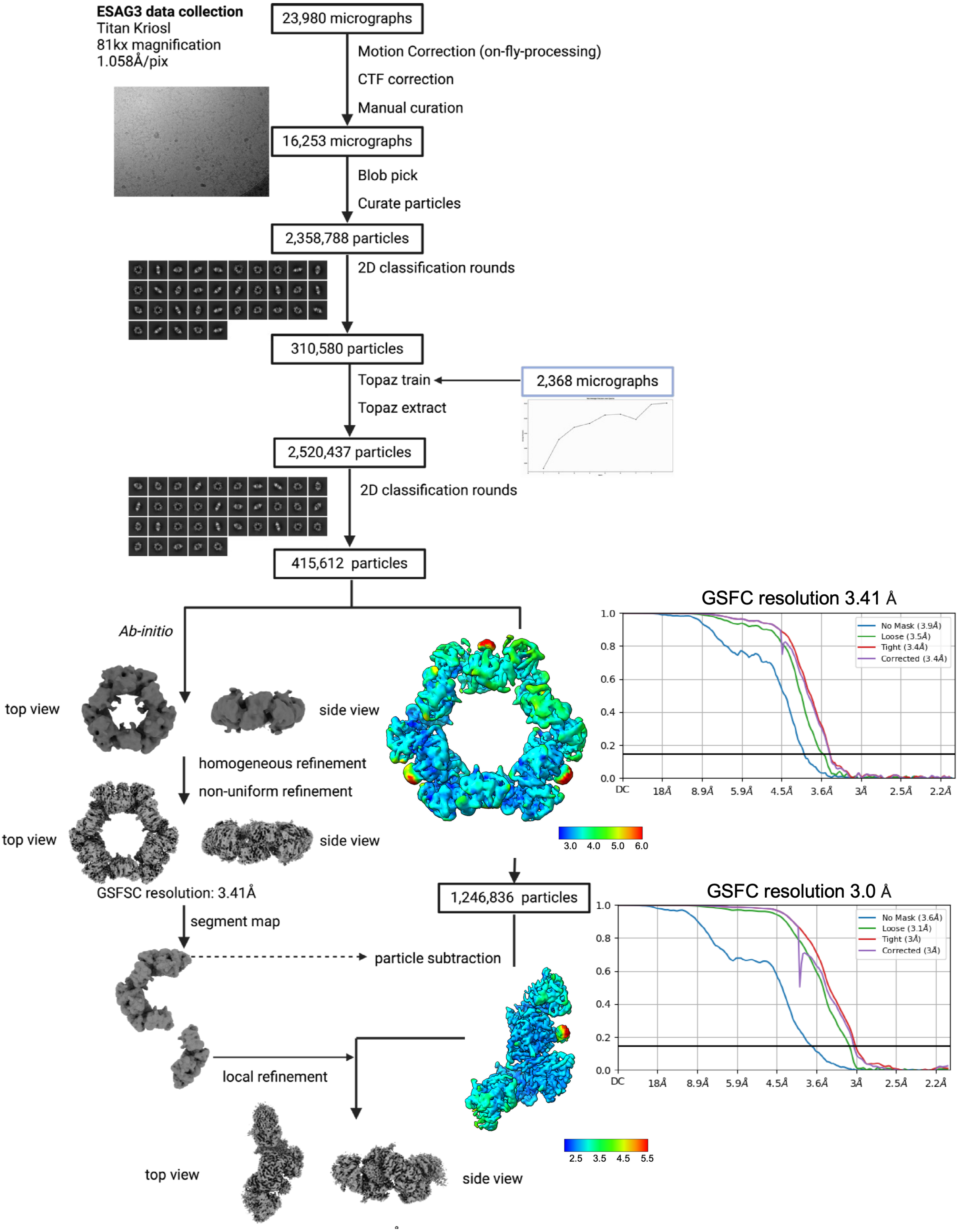


**Figure S4: CryoSPARC (v4) cryo electron microscopy data processing workflow.**
Cryo-EM data processing workflow for ESAG3. Micrographs were collected using a Titan Krios (X-FEG, K3, 300kV) at a dose of 39.94 e⁻/Å2 and 81kx magnification with a pixel size of 1.058 Å/pix. A total of 23,980

micrographs underwent motion correction (on-fly processing), CTF correction, and manual curation, yielding 16,253 micrographs. Following blob picking and particle curation, 2,358,788 particles were extracted and subjected to 2D classification rounds, resulting in 310,580 particles. Topaz training was performed using 2,368 micrographs, followed by Topaz extraction yielding 2,520,437 particles. Subsequent 2D classification rounds reduced the dataset to 415,612 particles. Ab-initio reconstruction (top and side views shown) was performed, followed by homogeneous refinement, which aligns all particles to a single structure, and non-uniform refinement, which enhances resolution in rigid regions while maintaining flexibility in others, yielding a ring structure with a GSFC resolution of 3.57 Å (coloured structure and FSC curve shown). C3 symmetry expansion was applied, followed by particle subtraction of two-thirds of the ring lobes using a mask generated in ChimeraX (v1.8). The resulting 1,246,836 particles were used for local refinement of a single subunit, achieving a final resolution of 3.00 Å (coloured subunit structure and FSC curve shown). Grey structures represent intermediate reconstruction models (top and side views). Colour gradient represents local resolution range (2.5 to 6.0 Å) in both hexamer and octadecamer.

#### Table S2 Data collection and refinement statistics

| Parameter | ESAG3 ring (homo-octadecamer) | ESAG3 subunit (hexamer) |
|---|---|---|
| **Data collection** | | |
| Facility | eBIC / Diamond Light Source | |
| Microscope | Titan Krios m07 | |
| Detector | Gatan K3 | |
| Electron source | Schottky X-FEG | |
| Voltage (kV) | 300 | |
| Electron exposure ($e^{-}$/Å$^2$) | 39.94 | |
| Magnification | 81,000× | |
| Pixel size (Å) | 1.058 | |
| Defocus range (µm) | 0.1-4.0 | |
| --- | --- | --- |
| **Data processing** | | |
| Software | **cryoSPARC v4, Phenix, ISOLDE, CCPEM** | |
| Micrographs collected (no.) | 23,980 | |
| Micrographs used (no.) | 16,253 | |
| Initial particle images (no.) | 2,520,437 | |
| Final particle images (no.) | 415,612 | 1,246,836 |
| Symmetry imposed | C3 | C1 (local refinement) |
| --- | --- | --- |
| **Refinement** | | |
| Map resolution (Å) | 3.4 | 3 |
| FSC Threshold | 0.143 | 0.143 |
| Map sharpening B-factor (Å$^2$) | -157.6 | -109.6 |
| Initial model used | **AlphaFold3 ESAG3 monomer, docked in UCSF ChimeraX, refined in ISOLDE** | |
| Model composition | | |
| Protein residues | 5904 | 1968 |
| Non-hydrogen Atoms | 48330 | 16110 |
| B-factors (Å$^2$) | 235.4 Å$^2$ | 200.1 Å$^2$ |
| Protein | | |
| R.m.s. Deviations | | |
| Bond lengths (Å) | 0.01 | 0.01 |
| Bond angles (°) | 1.87 | 1.79 |
| --- | --- | --- |

#### Figure S5

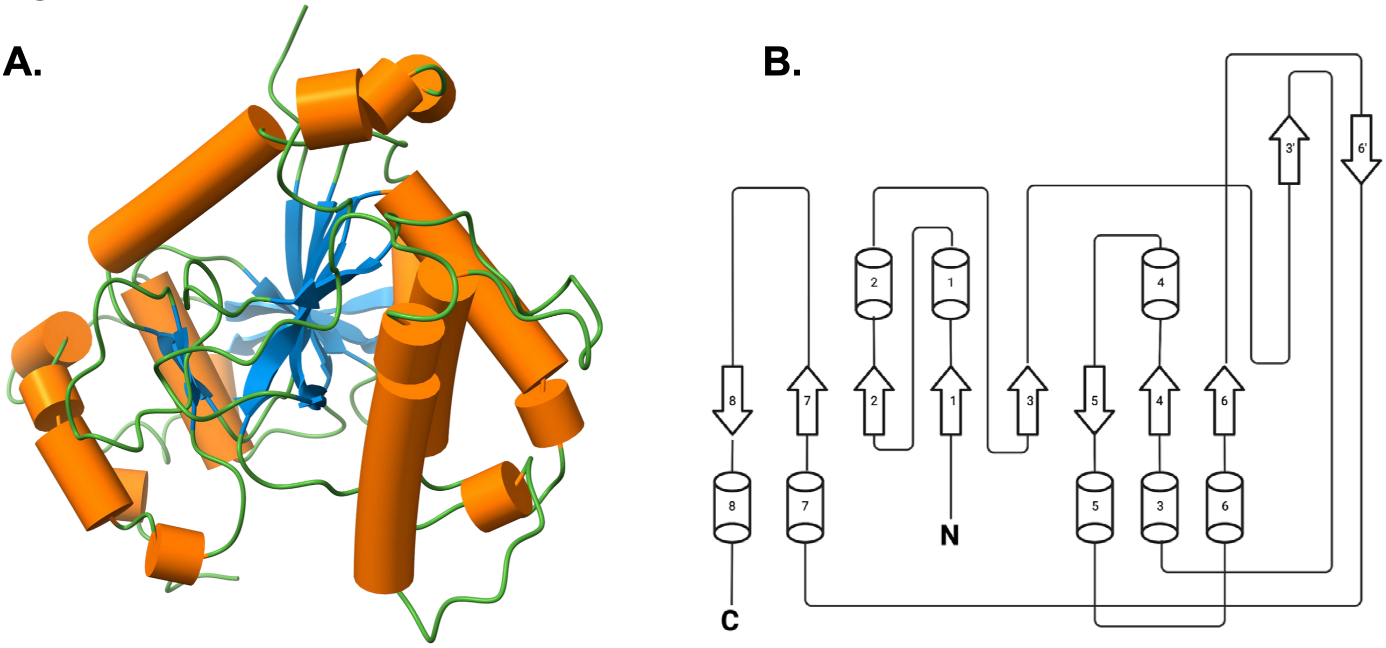


**Figure S5. ESAG3 adopts a glycosyltransferase-A (GT-A) family fold**

**A. Cartoon representation of an ESAG3 protomer.** The structure displays the characteristic GT-A Rossmann-like α/β/α sandwich architecture with alternating α-helices (cylinders) and β-sheets (arrows).

**B. Topology diagram illustrating the ESAG3 secondary structure arrangement.** Cylinders indicate α-helices; arrows indicate β-strands. The conserved GT-A architecture features a central β-sheet core flanked by α-helices. N- and C-termini are labelled.

**Figure S6**

**A.**

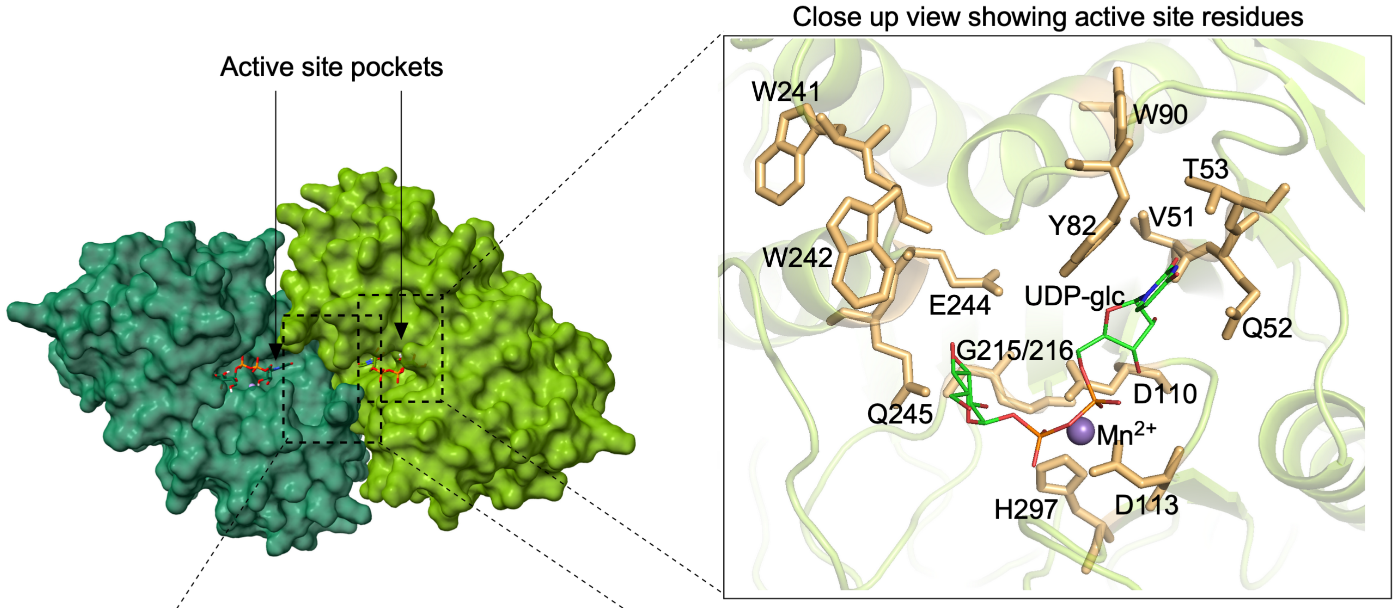
Active site pockets
Close up view showing active site residues
W241
W90
T53
Y82
V51
W242
E244
UDP-glc
Q52
G215/216
D110
Q245
Mn2+
H297
D113

**B.**

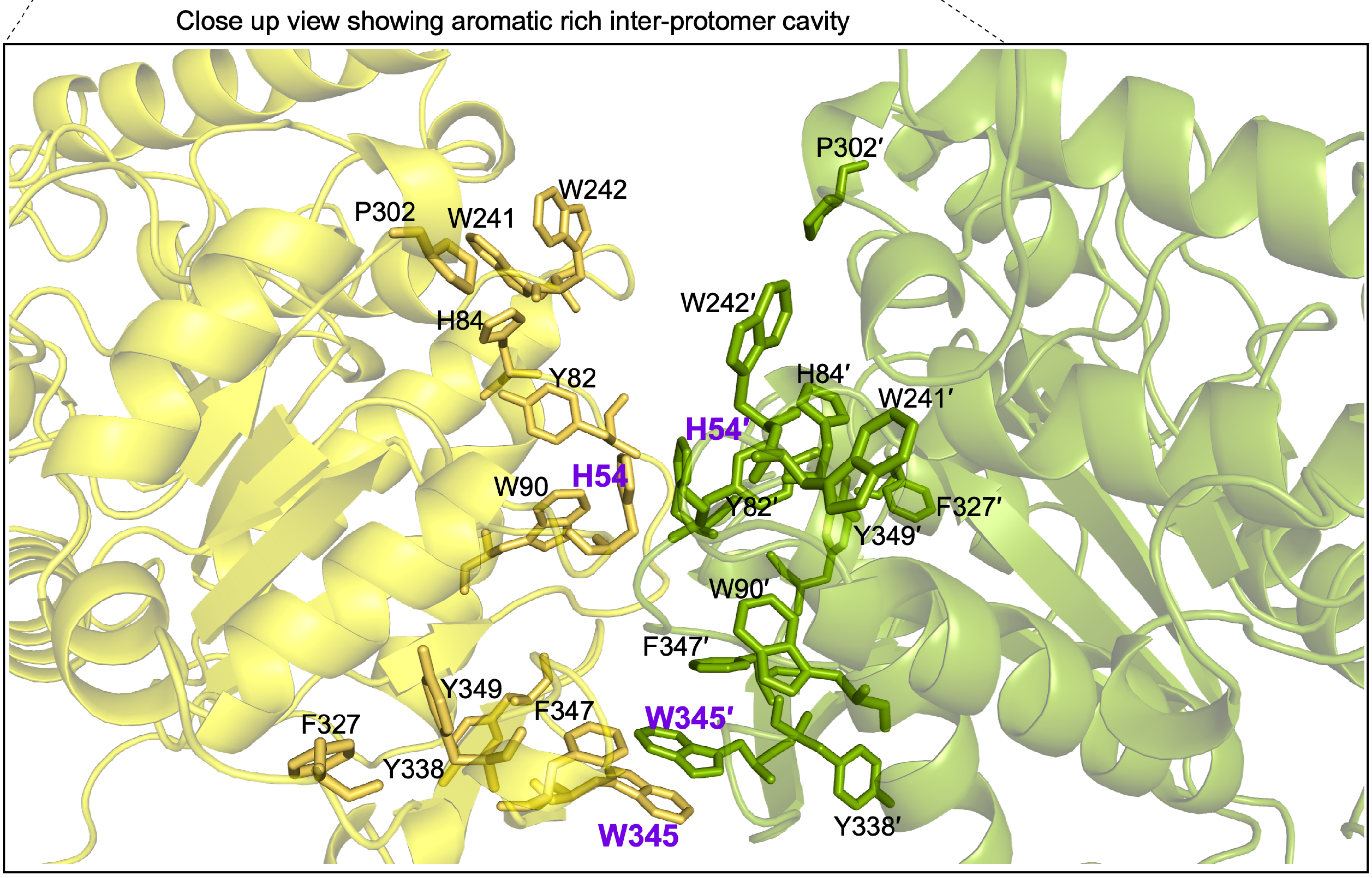
Close up view showing aromatic rich inter-protomer cavity
P302′
W242
P302
W241
W242′
H84
H84′
Y82
W241′
H54′
W90
H54
Y82′
F327′
Y349′
W90′
F347′
Y349
F347
F327
W345′
Y338
W345
Y338′

**Figure S6. Architecture of ESAG3 active site and dimer interface showing aromatic rich contacts between protomers**

**A. Surface view of a head-to-head ESAG3 dimer (teal/lime). Left:** The individual GT-A active-site clefts from each protomer face inward, creating a single bilobed cavity between protomers. Arrows indicate the positions of modelled UDP donor sites transplanted from the LH3 structure. **Right**: Close-up of one protomer active site with UDP-Glc and $Mn^{2+}$ positioned by superposition with LH3 (PDB: 6FXR). Residues forming the donor-site rim are labelled: V51, Q52, T53, H54, Y82, W90, E244/Q245 (XED region), G215–G216 (glycine loop), D110/D113 (DxxD metal loop), H297, W241 and W242. This panel illustrates all residues important for catalysis are contained in one protomer.

**C. Close-up of the ESAG3 dimer interface.** Head-to-head protomers (yellow/green) interlock across an aromatic-rich pocket adjacent to each active site. Interface residues include H54, Y82, H84, W90, W241, W242 (N-patch) and F327, Y338, W345, F347, Y349 (C-patch); primes denote the partner protomer. ChimeraX contact analysis predicts π–π stacking for H54/H54′ and W345/W345′. Mutated positions H54 and W345 used in Fig. 4 are highlighted in purple.

#### Figure S7

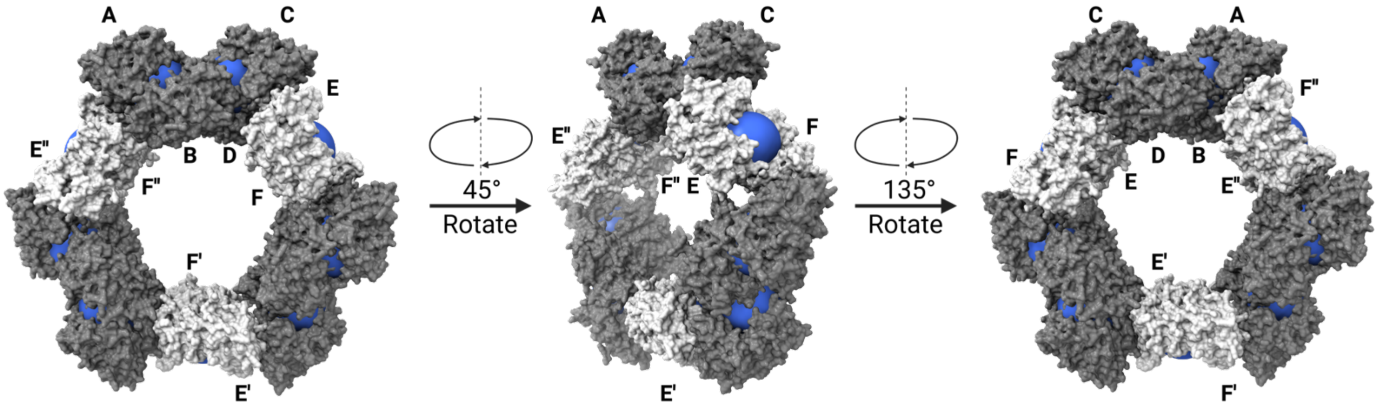


**Figure S7. Orientation and solvent accessibility of inter-protomer cavities in the ESAG3 octadecameric ring.**
Surface representations of the ESAG3 octadecamer are shown at three viewing angles, obtained by successive rotations of 45° and 135° around the ring axis. Three outward-facing head-to-head dimers (EF, E′F′ and E″F″) are highlighted in light grey, while dimers whose inter-protomer cavities are oriented away from the ring periphery and are partially occluded by neighbouring protomers are shown in dark grey.

Blue spheres indicate the approximate location and extent of the enlarged inter-protomer cavity formed at each head-to-head dimer interface. For each dimer, the sphere was centred at the geometric centre of the catalytic DXXD motif (Asp110 and Asp113) and the C-terminal His297 from both protomers, and rendered with a radius of 18 Å. Among the nine dimers in the octadecameric assembly, three (shown in light grey) present outward-facing, solvent-accessible inter-protomer cavities, while the remaining six (shown in dark grey) contain cavities that are partially occluded by neighbouring protomers.

#### Figure S8

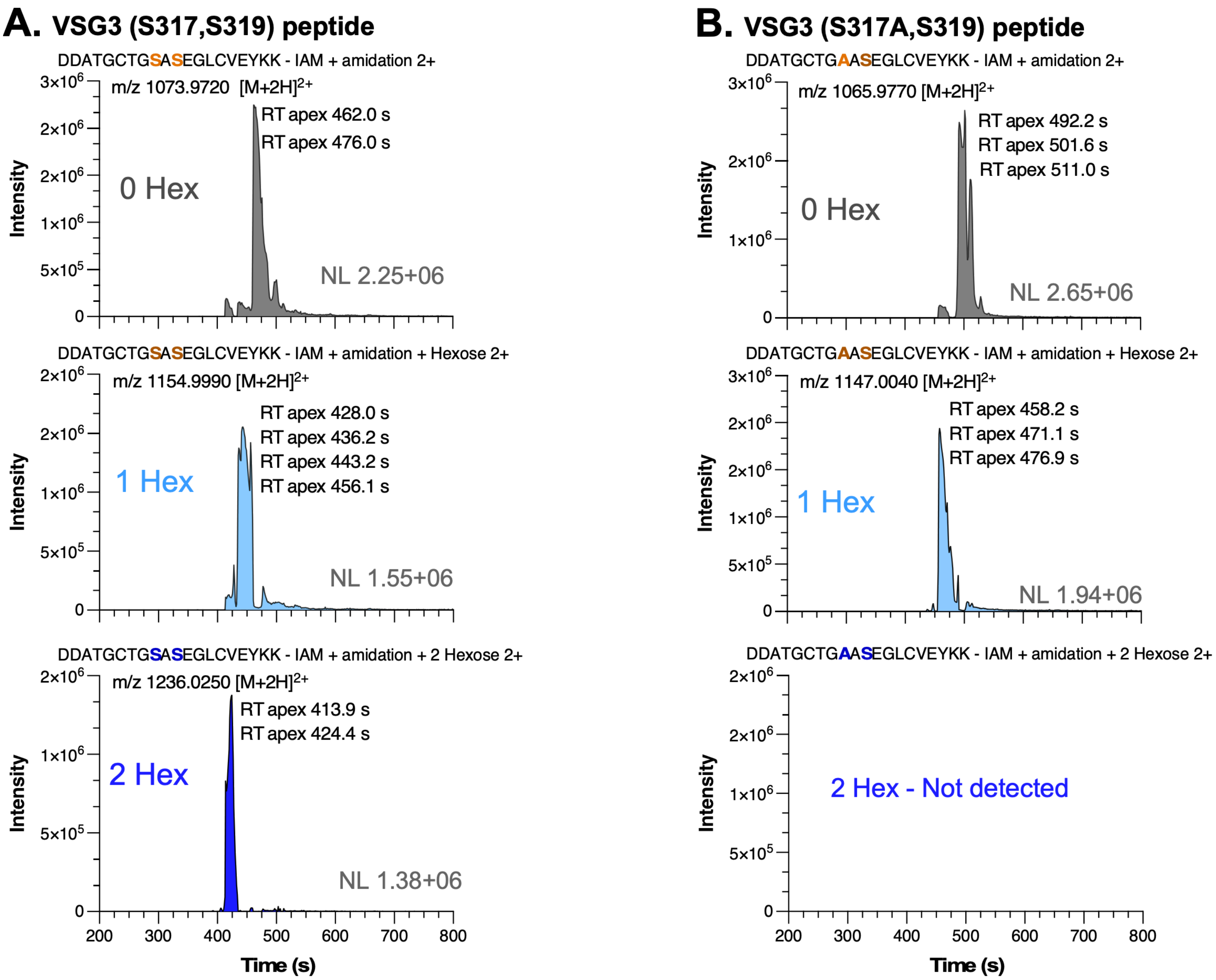


**Figure S8.** Accurate-mass extracted ion chromatograms of VSG3 peptide glucosylation reactions by H54A/W345A ESAG3.

Extracted ion chromatograms for the $[M+2H]^{2+}$ ions of unmodified (0 Hex), monohexosylated (1 Hex), and dihexosylated (2 Hex) VSG3 peptides following incubation with H54A/W345A ESAG3 from SEC Peak fraction 2 in **Fig 5**. In **A and B**, we refer to hexosylated products as glucosylated since only UDP-glucose were include in reaction mixtures.

**A** VSG3(S317,S319) peptide incubated with H54A/W345A ESAG3. Extracted ions correspond to unmodified peptide (m/z 1073.9720), monoglucosylated peptide (m/z 1154.9990) and diglucosylated peptide (m/z 1236.0250).

**B** VSG3(S317A,S319) peptide incubated with H54A/W345A ESAG3. Extracted ions correspond to unmodified peptide (m/z 1065.9770) and monoglucosylated peptide (m/z 1147.0040). Diglucosylated product was not detected.

Each trace is autoscaled to its own maximum; **NL** indicates the absolute maximum intensity (detector counts). Retention times of resolved features are annotated. No dihexosylated product is detected on the S317A, S319 VSG3 peptide in (**B**). Integrated peak areas for all glycoforms (combining $[M+2H]^{2+}$ and $[M+3H]^{3+}$) and the calculated percentages correspond to the relative glycoform distributions shown in **Fig. 5**.

#### Figure S9

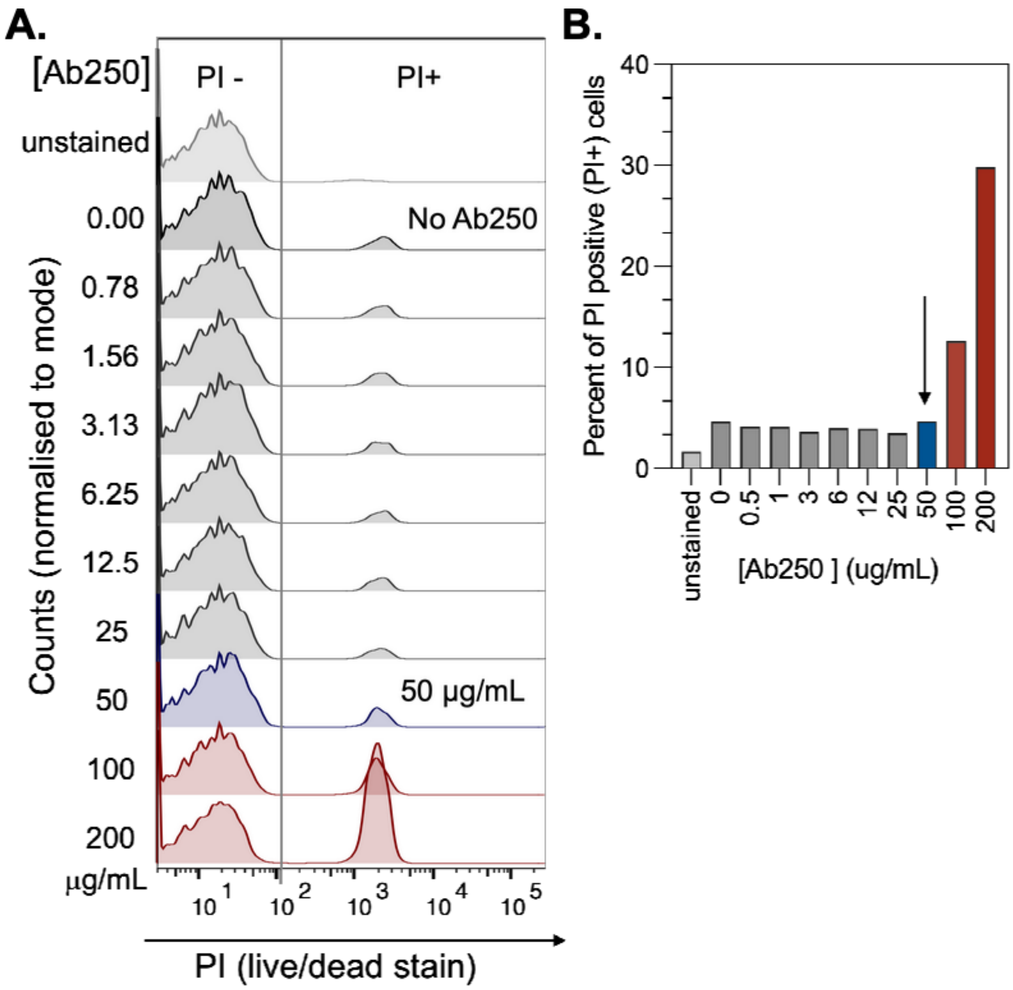


**Fig S9: Determination of effective Ab250 concentration that minimises antibody-mediated cytotoxicity in VSG3$^{WT}$ expressing trypanosomes independent of complement.**

**A.** Bloodstream-form trypanosomes expressing VSG3-WT were incubated with a 1:2 serial dilution series of Ab250 (200–0.5 µg/mL) for 10 min at 4°C, followed by 30 min at 37°C in the absence of complement. A no-antibody PI-stained control and an unstained baseline control were included. Cell viability was assessed by propidium iodide (PI) exclusion by flow cytometry, gated on single trypanosomes (minimum 6,000 events per sample). Representative histograms are shown; n = 1 biological replicate. Technical duplicates for 200, 100, and 50 µg/mL are not shown but consistent with above result.

**B.** Quantification of percentage PI-positive cells across the dilution series. Antibody-dependent cytotoxicity, independent of complement, was observed at 200 µg/mL (~30% PI-positive) and 100 µg/mL (~12% PI-positive). Cell viability at 50 µg/mL and below was indistinguishable from the no-antibody control (~1.4–2% PI-positive). Arrowhead indicates 50 µg/mL, selected as the working concentration for subsequent complement killing experiments to avoid confounding complement-independent cytotoxicity effects.
